# METRIN-KG: A knowledge graph integrating plant metabolites, traits, and biotic interactions

**DOI:** 10.1101/2025.08.20.671289

**Authors:** Disha Tandon, Tarcisio Mendes De Farias, Pierre-Marie Allard, Emmanuel Defossez

## Abstract

**Background:** In recent years, biodiversity data management has emerged as a critical pillar in global conservation efforts. Today, the ability to efficiently collect, structure, and analyze biodiversity data is central to breakthroughs in conservation, drug development, disease monitoring, ecological forecasting, and agri-tech innovation. However, due to the vastness and heterogeneity of biodiversity data, it is often confined to databases for specific research areas in isolated formats and disconnected from other relevant resources. Crucial components of such data in kingdom Plantae comprise of metabolomes - the vast array of compounds produced by plants; traits - measurable characteristics of plants that influence their growth, survival, and reproduction, and that affect ecosystem processes; and biotic interactions - relationships of plants with other living organisms, affecting the ecosystem functions.

**Results:** In this work, we present METRIN-KG (MEtabolomes, TRaits, and INteractions-Knowledge Graph) a powerful data resource simplifying the integration of diverse and heterogeneous data resources such as plant metabolomes, traits, and biotic interactions.

**Conclusions:** The proposed knowledge graph provides an interface to interactively search for data relating plant metabolomes, traits, and interactions. This, in turn, will facilitate development of research questions in life-sciences. In this context, we provide representative case studies on how to frame queries that can be used to search for relevant data in the knowledge graph.

## Context

### Interplay of metabolomes, traits, and biotic interactions

All species of our planet are connected by means of shared resources for sustenance, response to environmental effects, and multi-level biotic and abiotic interactions. An in-depth understanding of this multi-scale network of interactions enables researchers from natural sciences, chemistry, microbiology, ecology, plant biology, and climate change to address critical questions in ecology, biodiversity conservation, agriculture, and human health. As referenced by studies before [1–3], there are seven main shortfalls of biodiversity knowledge. Amongst these shortfalls, Raunkiæran and Eltonian emphasize *lack of connected knowledge on species’ traits and interactions, as well as their corresponding relation to ecological functions*. The chemistry of life, by governing biotic interactions, resource access, environmental adaptation, and individual phenotype, provides a link to building the mechanistic network required to understand species’ processes underlying ecosystem functioning [4].

The metabolome, a key component of the chemistry of life, refers to all metabolites forming the substrates or products of enzymatic reactions in an organism. To a certain extent, the metabolome provides a bridge between the ecosystem functioning and the contextual information of species’ states across multiple scales (spatial, temporal, and environmental) [5–7]. In the Plantae kingdom, the sheer magnitude of metabolite diversity produced by plants is estimated to be between 1.5 and 25.7 million collectively for 400,000 plants [8], which poses significant challenges for metabolomic data analysis [9,10].

It has been shown that metabolomes can provide a proxy for the estimation of the plant functions, thus revealing the cause-and-effect relationship between external environmental factors and plant fitness [9]. However, exploring the metabolome structure across spatial, temporal, and environmental scales requires extensive studies to unravel the interplay of hundreds of compound classes and their potential biological and ecological functions.

Since experimental approaches alone cannot provide such comprehensive datasets, aggregating data across multiple studies that combine metabolome with physiological or environmental data appears as a necessary solution. Such information can include functional data like traits and co-dependence data like interactions.

Plant ecology focuses on how an organism’s traits influence its interactions with and in response to environmental conditions throughout its life cycle. Traits like plant height, seed mass, leaf area, leaf carbon, nitrogen, and phosphorus contents have been used to define plant processes across taxonomic (tree of life) [11–13], spatial (e.g.: subalpine regions) [14], environmental (e.g.: weather changes) [15], and temporal (e.g.: reproduction during the life of plant or life history) scales [15–17]. However, the mechanistic understanding behind the effect of traits on plant fitness and functioning is not clear [18–21] because of the unexplained or ambiguous relation of traits and variation in ecosystem functioning.

Plant biotic interactions, including relationships with other plants, fungi, bacteria, and soil organisms, have been shown to influence ecosystem structure and function, ultimately shaping biodiversity patterns [22–24]. As sessile organisms, plant biotic interactions also rely heavily on chemical mediation, involving both volatile and non-volatile compounds to compensate for their lack of mobility. Such interactions, particularly those involving insects, such as defense mechanisms or pollinator attraction, have been closely linked to the diversity of plant metabolites through evolutionary processes [25–29].

### Challenges in integrating high-dimensional data

In recent years, efforts have been made to include chemistry into the functional traits framework to resolve plant functions in ecosystems [30,31]. Many studies have tried to combine the traits with specific compounds or classes of compounds [30–34], biotic interactions with traits [35–38], and biotic interactions with metabolomes [39–42]. Some studies have combined the three to decipher ecosystem functioning. For instance, they have examined plant traits, their interactions with soil biota, and related chemodiversity [43,44]. Some have explored the effect of insect herbivory on plant traits and their secondary metabolite concentrations [45–47]. Others investigated links between plant root traits, nutrient foraging, chemodiversity, and their symbiotic relationship with mycorrhizal fungi [48–50]. Additional research has focussed on allelopathic interactions and nutrient mobilization during intercropping in agriculture [51,52]. Studies also cover plant root traits, exudates (metabolites produced by roots), and its interactions with the rhizosphere microbiome [53]. Further examples include links between biotic interactions, elevation gradients, and metabolomics [54]; climate-induced plant host shift in insects [55]; plant microbe interactions and chemical defense mechanisms [56]; as well as insect herbivory and defense mechanisms facilitated by secondary metabolites [57–59]. Moreover, several reviews published during the last decade have stressed the importance of combining plant traits, interactions, and metabolomes [33,60–67]. The main challenge in studies that combine these three components is the high dimensionality of each, coupled with limited resources available to characterize such complexity, which poses analytical limitations (e.g. incomplete spectral libraries, detection limits), suboptimal data availability, and cross-links across the three components.

The experimental approaches commonly used in functional ecology are powerful tools for exploring and describing specific mechanisms. However, they remain limited when it comes to disentangling complex processes operating at larger scales. Linking chemical pathways, compound classes, or molecular structures to ecological functions remains a major challenge in ecological research. Recent development of databases and advances in data science offer a new potential to reach this goal. A data-driven approach is required to first map the existing knowledge in all three areas followed by the development of research hypotheses. Building on the few previous studies in this direction, we explored various datasets and databases related to traits, interactions, and metabolomics, which ultimately inspired the core idea of this study.

Much information on the heterogeneity of metabolites biosynthesized by organisms lies locked in isolated tabular formats, excel sheets, and pdf documents (as discussed in [68]). Few databases provide comprehensive information on metabolites or natural products [68–72]. The Earth Metabolome Initiative (EMI) [73] and its pilot project Digital Botanical Gardens Initiative (DBGI) [74], were launched in 2022. They aim to document metabolic content for all known species on Earth (initiating point being botanical gardens living collections), following the Findable, Accessible, Interoperable, Reusable (FAIR) guidelines [75]. Under the EMI umbrella, comes the Experimental Natural Products Knowledge Graph (ENPKG) [69], a published resource of metabolomes from 1600 tropical plant extracts. ENPKG uses a sample-centric approach comprising semantic annotations to structure large, heterogeneous metabolomics datasets into knowledge graphs. It also enables harmonization of experimental data with publicly available resources. However, no ecological metadata was integrated in ENPKG.

Like metabolite data, multi-species level interaction data also lies locked as pairwise correlation metrics in peer-reviewed research papers. While Global Biotic Interactions (GloBI) [76] and tools like BiotXplorer [77] have succeeded in collecting this information in a FAIR resource [78], they allow only partial multi-species level interaction mapping and are limited to pairwise interactions. Such documentation of interactions is remarkable, yet it suffers from the lack of providing complete multi-species level interactions. Furthermore, higher-order interaction structures (e.g. multi-trophic or trait-mediated interactions) are often analyzed post hoc using correlation-based summaries that abstract away mechanistic or phenotypic information. Similarly, data on plant trait heterogeneity is limited to few resources like TRY [79,80] and global plant trait network [81].

There are individual databases listing organism traits [80,82], metabolite-pathway relations of a limited number of plant species [83] as well as microbes [84,85], pairwise interaction maps [76], medicinal herbs [86], and food [87]. However, there is no single resource available that combines knowledge across organisms, detailing their traits, interactions, and their complete/partial metabolomes, all of which are important for the in-depth understanding of the complex network of life.

### MEtabolomes, TRaits, and INteractions - Knowledge Graph (METRIN-KG)

In this Data Note, we present the first efforts in combining publicly available data on plant traits, interactions, and metabolome from peer-reviewed research and databases in METRIN-KG. We have linked enriched metabolome datasets behind ENPKG [69] with high dimensional data on plant traits from TRY database [80], pairwise interaction data from GloBI [76], and annotated data on natural products from LOTUS [68] (available through Wikidata). To ensure semantic interoperability, we have used ontologies for knowledge representation like the Earth Metabolome Initiative Ontology (EMI Ontology) [88,89], and the ENPKG ontology for natural product-specific concepts [69]. We used Ontop [90–92] and Python rdflib library [93] to materialize the Resource Description Framework (RDF) triples. We further implemented a SPARQL Protocol and RDF Query Language (SPARQL) editor to query METRIN-KG and retrieve results.

We anticipate METRIN-KG to be useful for interactively searching related information on traits, interactions, and metabolome, thus guiding and inspiring the development of future research questions in the fields of ecology, biodiversity conservation, agriculture and human health. Moreover, we believe this information could also be valuable to a wider audience beyond researchers, such as policymakers and public health professionals, by enhancing their understanding of the broader implications of research within their respective fields.

In the following sections we provide details on how the datasets used in METRIN-KG were retrieved, structured, and linked. We also provide a brief overview of the construction of the EMI Ontology, discuss potential reuse of METRIN-KG, and present representative queries for exploring the knowledge graph.

## Methods

### Data retrieval from TRY database

For this study, the pilot dataset was retrieved from the TRY website [94] by requesting data for 41 traits based on the most used functional trait categories in plant physiology studies [30] - plant height, seed mass, leaf area, leaf carbon content, leaf nitrogen content, leaf phosphorus content, stem specific density, leaf lifespan, leaf respiration rate, and photosynthesis rate (**Table-1**). Scientific names of the species from this dataset were mapped to Wikidata as listed in the section ‘Taxonomy mapping’. The retrieved data is archived publicly in file ‘TRYdb_40340.txt.gz’ in a Zenodo repository [95].

**Table-1:** List of traits used for retrieving data from TRY database.

| Trait identifier<br>in TRY | Trait Name |
| --- | --- |
| 1 | Leaf area |
| 3108 | Leaf area (in case of compound leaves: leaf, petiole excluded) |
| 3109 | Leaf area (in case of compound leaves: leaflet, petiole excluded) |
| 3110 | Leaf area (in case of compound leaves: leaf, petiole included) |
| 3111 | Leaf area (in case of compound leaves: leaflet, petiole included) |
| 3112 | Leaf area (in case of compound leaves: leaf, undefined if petiole in- or excluded) |
| 3113 | Leaf area (in case of compound leaves: leaflet, undefined if petiole is in- or excluded) |
| 3114 | Leaf area (in case of compound leaves undefined if leaf or leaflet, undefined if petiole is in- or excluded) |
| 11 | Leaf area per leaf dry mass (specific leaf area, SLA or 1/LMA) |
| 3115 | Leaf area per leaf dry mass (specific leaf area, SLA or 1/LMA): petiole excluded |
| 3116 | Leaf area per leaf dry mass (specific leaf area, SLA or 1/LMA): petiole included |
| 3117 | Leaf area per leaf dry mass (specific leaf area, SLA or 1/LMA): undefined if petiole is in- or excluded) |
| 3085 | Leaf area per leaf dry mass (specific leaf area, SLA or 1/LMA) of leaf lamina |
| 3086 | Leaf area per leaf dry mass (specific leaf area, SLA or 1/LMA) petiole, rhachis, and midrib excluded |
| 3106 | Plant height vegetative |
| 3107 | Plant height generative |
| 26 | Seed dry mass |
| 3660 | Seed fresh mass |
| 3836 | Seed mass per fruit |
| 4 | Stem specific density (SSD, stem dry mass per stem fresh volume) or wood density |
| 3452 | Stem specific density (SSD, stem dry mass per stem fresh volume) or wood density: heartwood |
| 3764 | Stem specific density (SSD, stem dry mass per stem fresh volume) or wood density: stem |
| 13 | Leaf carbon (C) content per leaf dry mass |
| 14 | Leaf nitrogen (N) content per leaf dry mass |
| 15 | Leaf phosphorus (P) content per leaf dry mass |
| 12 | Leaf lifespan (longevity) |
| 663 | Leaf respiration rate in light per dry mass |
| 512 | Leaf respiration rate in light per leaf area |
| 664 | Leaf respiration rate in light per leaf respiration rate in the dark |
| 2943 | Leaf respiration rate in the dark as a fraction of photosynthetic carboxylation capacity (V<sub>c</sub>max) |
| 665 | Leaf respiration rate in the dark minus respiration in light, mass based |
| 54 | Leaf respiration rate in the dark per leaf area |
| 41 | Leaf respiration rate in the dark per leaf dry mass |
| 500 | Leaf respiration rate in the dark per leaf nitrogen (N) content |
| 272 | Leaf respiration rate in the dark temperature dependence |
| 53 | Photosynthesis rate per leaf area |
| 974 | Photosynthesis rate per leaf area: transition to TPU limited photosynthesis |
| 40 | Photosynthesis rate per leaf dry mass |
| 135 | Photosynthesis rate per leaf nitrogen (N) content (photosynthetic nitrogen use efficiency, PNUE) |
| 134 | Photosynthesis rate per leaf transpiration (photosynthetic water use efficiency: WUE) |
| 3128 | Photosynthesis rate per stomatal conductance |

*Abbreviations* - SLA: Specific Leaf Area; LMA: Leaf dry Mass per Area; SSD: Stem Specific Density; TPU: Triose Phosphate Utilisation; PNUE: Photosynthetic Nitrogen Use Efficiency; WUE: Water Use Efficiency

### Data retrieval from GloBI

The compressed stable release of interaction data was downloaded from the Zenodo archive of GloBI, version 0.8 from January 2025 [96], representing ‘species interactions tabulated as pairwise interactions in a zipped tab-separated values format.’ (as described on the GloBI website [97]). The creators of GloBI mention that ‘included taxonomic names are not interpreted, but included as documented in their sources’. The taxonomic identifiers and names were mapped to Wikidata as listed in the section ‘Taxonomy mapping’. The data is available publicly in file ‘verbatim-interactions.tsv.gz’ at GloBI Zenodo repository [96].

### Taxonomy mapping

GloBI collates interaction information from standardized data from numerous sources including online repositories and projects (e.g.: Encyclopedia of Life) as well as data entered by individual projects through GloBI’s dataset-template [98]. The full list of data sources is available on its website [99]. The combination of these two approaches resulted in taxonomic identifiers from around 15 taxonomy databases (**Supplementary Table 1**) to be present in GloBI. The initial taxonomic mappings from these 15 resources as well as their scientific names to Wikidata identifiers were done using QLever’s [100] SPARQL user interface (UI) for Wikidata [101]. SPARQL queries used for mapping are as follows:

a) Query [102] for mapping Wikidata identifiers to ones from 15 taxonomy databases **(Supplementary Table 1).**
b) Query [103] for retrieving Wikidata identifiers and their lineage.

The full queries are listed in **Supplementary data.**

The retrieved data is available publicly in ‘wdTax’ files at METRIN-KG Zenodo repository [104].

For the TRY database, the scientific names were directly matched to ones obtained from Wikidata, to obtain corresponding Wikidata identifiers.

### Metadata mapping

GloBI provides limited mappings of life stages and body parts to Uber-Anatomy Ontology (UBERON) [105], Plant Ontology (PO) [106], Environment Ontology (ENVO) [107], Gene Ontology (GO; body parts only) [108], and Phenotype and Trait Ontology (PATO; body parts only) [109] for organisms involved in interactions (**Supplementary Table 2**). For biological sex, the names are provided in GloBI, but no specific mapping to existing ontologies.

With such limited mappings, it was crucial to map the raw text in metadata columns-body part, life stage, and biological sex. Such unmapped text was not single standardized terms, rather it contained compound expressions, abbreviations, counts, symbols, or multiple entities within a single field. For example, body part entries included values such as “2 guts”, “abdominal cavity”, “abo”, “abomasum/si”. Biological sex fields contained symbolic or aggregated values (e.g. “+”, “-”, “?”), numeric summaries (e.g. “10M,9F”, “11 males, 4 females, 2 juveniles”), or multilingual expressions (e.g. “11 machos | 9 hembras”). Life stage fields similarly contained compound or inconsistent representations, such as “adult; adult; egg”, “adult; larvae”, “0, I, II, III”, or mixed capitalization and punctuation.

As part of the metadata mapping process, such entries were first parsed using delimiters and normalization rules to separate them into individual candidate terms, which were then mapped independently to ontology entities.

To semantically match candidate terms with ontology concepts, we developed a script using ontology parsing and sentence embeddings. The code including the parsing and semantic matching [110] is publicly available at the METRIN-KG github repository. We utilized the Python Owlready2 library version-0.47 [111] to load a suite of biomedical and environmental ontologies, namely the ones originally present in GloBI, as well as others from their Persistent Uniform Resource Locators (PURL) (see **Supplementary Table 2**). For life stages and body parts, the full suite was considered, whereas for biological sex, only UBERON [105] and PATO [109] were considered.

From each ontology, we extracted class labels and associated synonyms (including exact, broad, and related synonyms where available) as candidate terms for matching. Synonyms were collected from Web Ontology Language (OWL) annotation properties such as ‘hasExactSynonym’, ‘hasBroadSynonym’, and ‘hasRelatedSynonym’. Each candidate term was paired with its primary label and class Internationalized Resource Identifier (IRI) for traceability.

To compute semantic similarity, we used the pre-trained ‘all-MiniLM-L6-v2’ model from Python SentenceTransformers library version-3.3.1 [112]. This model is a lightweight transformer trained to generate 384-dimensional dense vector embeddings that reflect the semantic content of short text spans. It is particularly suited for tasks such as semantic textual similarity and clustering.

All ontology terms (labels and synonyms) were encoded into dense vector representations using this model. Similarly, each user-provided input term for unmapped life stages, body parts, and biological sex was independently embedded into the same vector space. Term embeddings were computed using the model’s ‘encode()’ method with ‘convert_to_tensor=Truè, which produced PyTorch-compatible tensors suitable for high-performance vector operations.

Cosine similarity was computed between each input term vector and all ontology term vectors using the ‘util.pytorch_cos_sim’ function in SentenceTransformers. For each input term, the ontology term with the highest cosine similarity score was identified as the top candidate match. Cosine similarity is a common metric for comparing the orientation, rather than magnitude of two vectors. Given two vectors A and B, cosine similarity is defined as:

cosine_similarity(A,B)=A.B / |A|.|B|

Values range from -1 to 1, where 1 indicates identical direction (i.e., maximum similarity) and 0 indicates orthogonality (no similarity). For each input term, we computed its similarity against all ontology terms. The term with the highest similarity score was selected as the top candidate match. While all terms receive a candidate match, only matches with a similarity score ≥ 0.7 were flagged for manual review. This threshold ensures that low-confidence suggestions are carefully examined, while very low-similarity terms are treated as unmatched.

All results were output to a comma-separated values (CSV) file, including the input term, matched ontology label, its primary label, the ontology class IRI, and the similarity score.

While the automated process provided high-quality initial suggestions, all matches were manually reviewed and corrected to ensure semantic appropriateness. Corrections were informed by domain-specific knowledge and contextual relevance, particularly in cases where terms had multiple meanings or where high similarity scores did not guarantee ontological alignment. This two-stage process, automated semantic matching followed by manual curation, ensured both scalability and accuracy in aligning non-standardized input terms with controlled ontology concepts.

The above workflow was also run for matching the units of measure for trait data from TRY to the units vocabulary [113] provided by Quantities, Units, Dimensions, and Types (QUDT 2.1 schema) [114], followed by manual correction.

The mapped data is archived in the folders ‘globi’ and ‘trydb’ under processed data at METRIN-KG Zenodo repository [104].

### EMI Ontology

EMI [73] is a global effort to profile the metabolic content of all currently known species on our planet. Here, knowledge representation plays a key role to correctly capture, contextualize, and structure the vast amount of chemical diversity data that have been and will be generated in the upcoming years. Consequently, this will facilitate data (re)use and interoperability. To accurately represent the EMI knowledge, we explored several general-purpose and domain-specific ontologies to design a framework to describe chemical compounds (e.g., natural products) and their related data such as geolocation, provenance, organism sample metadata, and organism interactions. The EMI ontology reuses and repurposes, where applicable, several other ontologies beyond the life sciences such as World Wide Web Consortium’s (W3C) [115] Sensor, Observation, Sample, and Actuator (SOSA) ontology [116], that was primarily designed for other applications including the Open Geospatial Consortium (OGC) use cases. OGC is a consortium that aims to improve access to geospatial and location information [117]. Furthermore, semantic reconciliation powered by biocuration is at the core of our proposed ontology. For example, by applying the EMI ontology version 1.0, we can accommodate and harmonize different organismal, chemical and material sample taxonomies as well as vocabularies such as the Relation Ontology (RO) [118] to define interactions between organisms (e.g., “has pathogen”). The other ontologies reused in EMI are those of Simple Knowledge Organization System (SKOS) [119], standard units of measures from QUDT [114], geo-locations from the basic geo (World Geodetic System 1984 (WGS84) lat/long) vocabulary [120], and natural product-specific concepts from the ENPKG ontology [121]. Moreover, in addition to making several existing ontologies interoperable for the EMI knowledge representation, we created more than 100 new concepts and relations that can be easily identified since they start with the https://w3id.org/emi# (*emi:*) prefix in the EMI ontology [88], for example, the term *emi:NonTrait*, a non trait structured value. All new and reused terms that constitute the EMI ontology are documented and available online [88]. The code and a tutorial to build the metabolite-specific knowledge graph component of METRIN-KG with this ontology are available at the EMI Ontology GitHub repository [122]. **Figure-1** shows the EMI ontology schema.

**Figure-1:**
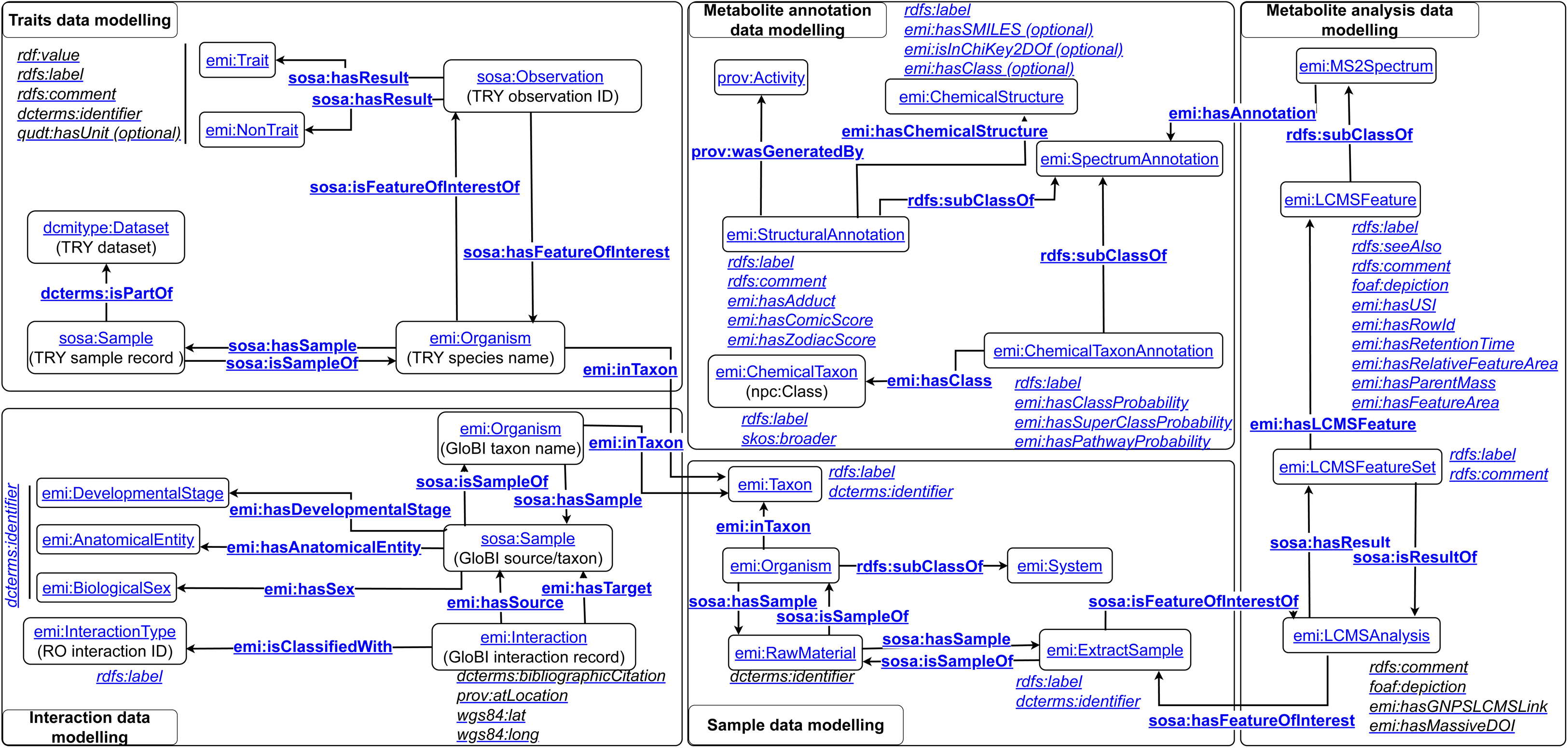
Snapshot of the data schema of the Earth Metabolome Initiative (EMI) ontology. Classes are represented in round-edged rectangles. Entity relationships are represented by directional arrows.

### Knowledge graph construction

Based on the EMI ontology as a data schema (**Figure-1**), we built a knowledge graph (KG) that semantically enriches, integrates, and interoperates 3 data sources - enriched metabolome datasets behind the ENPKG, trait data from TRY-db, and interaction data from GloBI. We developed the KG by combining subgraphs in two stages:

a) Ontop tool [90–92] was applied for developing the subgraph from enriched datasets (metabolite annotation, molecular networks, and taxonomical resolution results) originally used to build the ENPKG graph [123,124]. Ontop is a virtual knowledge graph system where SPARQL queries are translated into Structured Query Language (SQL) queries based on the predefined mappings. Ontop also provides means to materialize a KG according to these mappings. Therefore, to build the graph, one table for each tabular file was created in a relational database with a simplistic data schema. The files were loaded in this database through mysql version-8.2 [125]. Finally, to construct the KG, mappings were defined between the relational schema and the EMI ontology using the Ontop mapping language [90]. With these mappings [126], the EMI ontology [88], and the relational database, the RDF triples were materialized with Ontop to compose the KG.
b) For developing the subgraphs from TRY database and GloBI, Python rdflib library version-7.0.0 [93,127] was used. To construct them, the tab-separated value (TSV) files obtained for each of these two resources were used. These tables were then connected through Wikidata identifiers (**Figure-2**) for taxonomy, whenever they were available as indicated in section ‘Taxonomy mapping’. Mappings were defined between the relational schema and the EMI ontology using rdflib’s inherent capability to map ontology elements. The process for incorporating mappings to metadata for GloBI (source/target taxonomy, life stage renamed to developmental stage, body part renamed to anatomical entity, and biological sex) and TRY database (taxonomy and trait’s units of measure) was performed before generating the RDF triples. The code to develop the subgraphs is available at the METRIN-KG GitHub repository [128].

**Figure-2:**
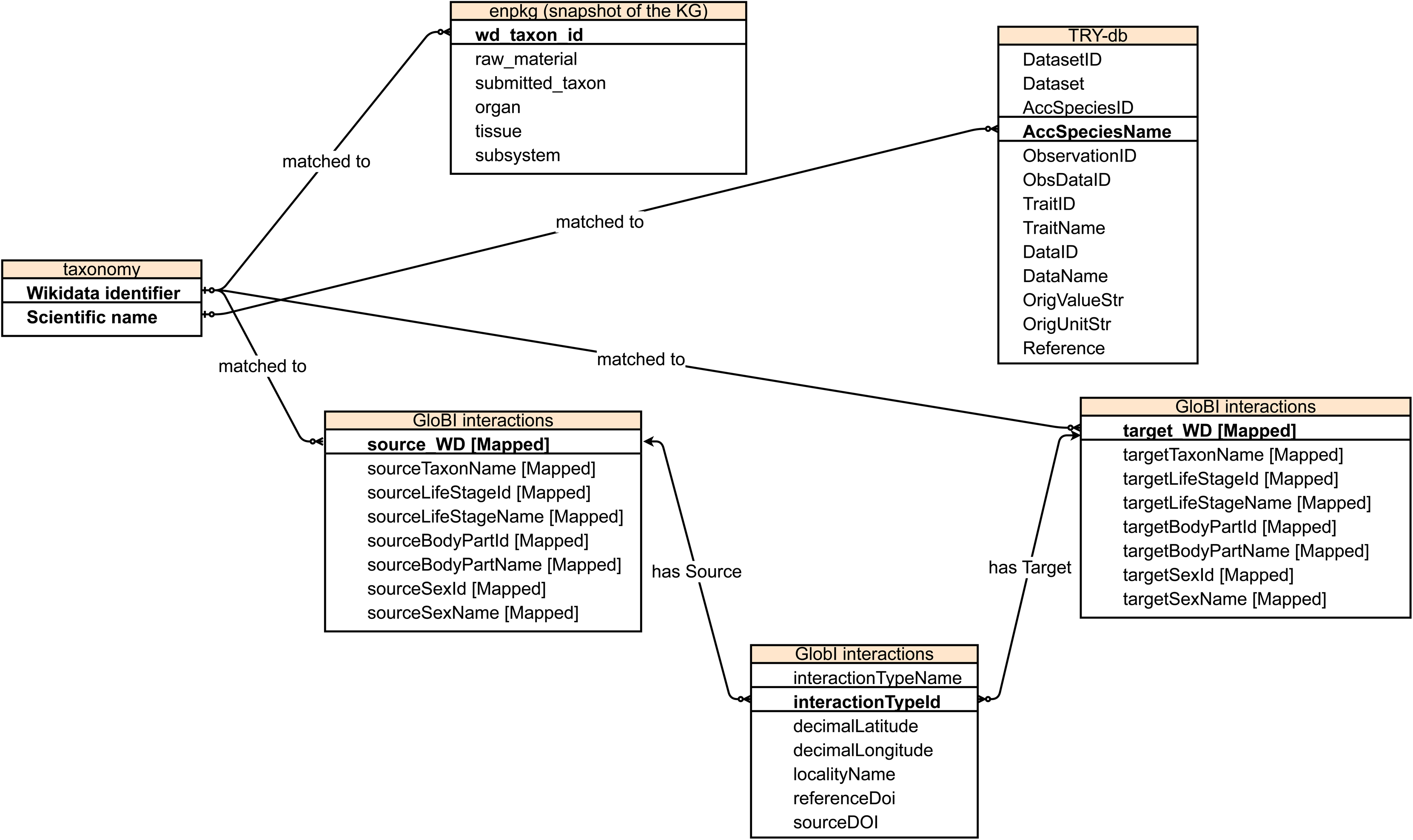
Schema of relations between different entities of integrated datasets of metabolome, traits, and interactions. Entities mapped are represented in bold and the corresponding relationship is represented by directional arrows.

The RDF files can be downloaded at METRIN-KG Zenodo repository [104] in the folder ‘KG’ under processed data.

### Indexing the knowledge graph and implementing the SPARQL end-point

We used Qlever [100], a SPARQL engine, to index our knowledge graph by implementing the steps described in qlever-control [129]. A SPARQL editor to query the indexed graph is available at [130]. This endpoint can also be accessed through https://kg.earthmetabolome.org/metrin/api [131] for programmatic purposes. An example for the programmatic access is provided on METRIN-KG’s github wiki [132]. The underlying operational endpoint is currently available at [133]. The stable URL [130], functions as a long-term and stable identifier to safeguard accessibility should the operational endpoint be modified in the future. This interface includes several examples covering specific use-cases for the three datasets (ENPKG metabolites, TRY-db traits, and GloBI interactions). Some of these use-cases are presented as case studies in the section ‘Data re-use and case studies’.

To provide the SPARQL editor user-interface for the indexed graph, we implemented qlever-ui [134,135] with few changes. To improve the user experience when navigating entities in the knowledge graph, we extended the qlever-ui interface to support fallback representations for non-available or missing web pages. This was achieved by incorporating SPARQL DESCRIBE queries into the frontend of the UI. The modified interface attempts to retrieve a minimal RDF-based summary of the requested entity when the original resource is not accessible (e.g., due to missing data in the knowledge graph). When such a condition is met, the interface automatically issues a SPARQL DESCRIBE query for the corresponding IRI. The result is rendered in a simplified RDF triple view, providing context about the entity based on available data in the backend knowledge base. The implementation is available via a fork of the official qlever-ui GitHub repository here [136].

## Data Description

### Mapping of TRY data

The original structure of the TRY datasets is modelled as plant species mapping to trait and non-trait data as well as related metadata like information on studies from which the data was obtained, as shown in **Figure-3**. The column names depicted in the figure are taken directly from the CSV table retrieved from TRY, the description of which is provided in **Supplementary Table 3**. Columns retained in the knowledge graph are indicated in the table and the figure. Overall, 20,272,589 records for 70,748 unique plant species were retrieved in a TSV format. (**Supplementary Table 4**).

**Figure-3:**
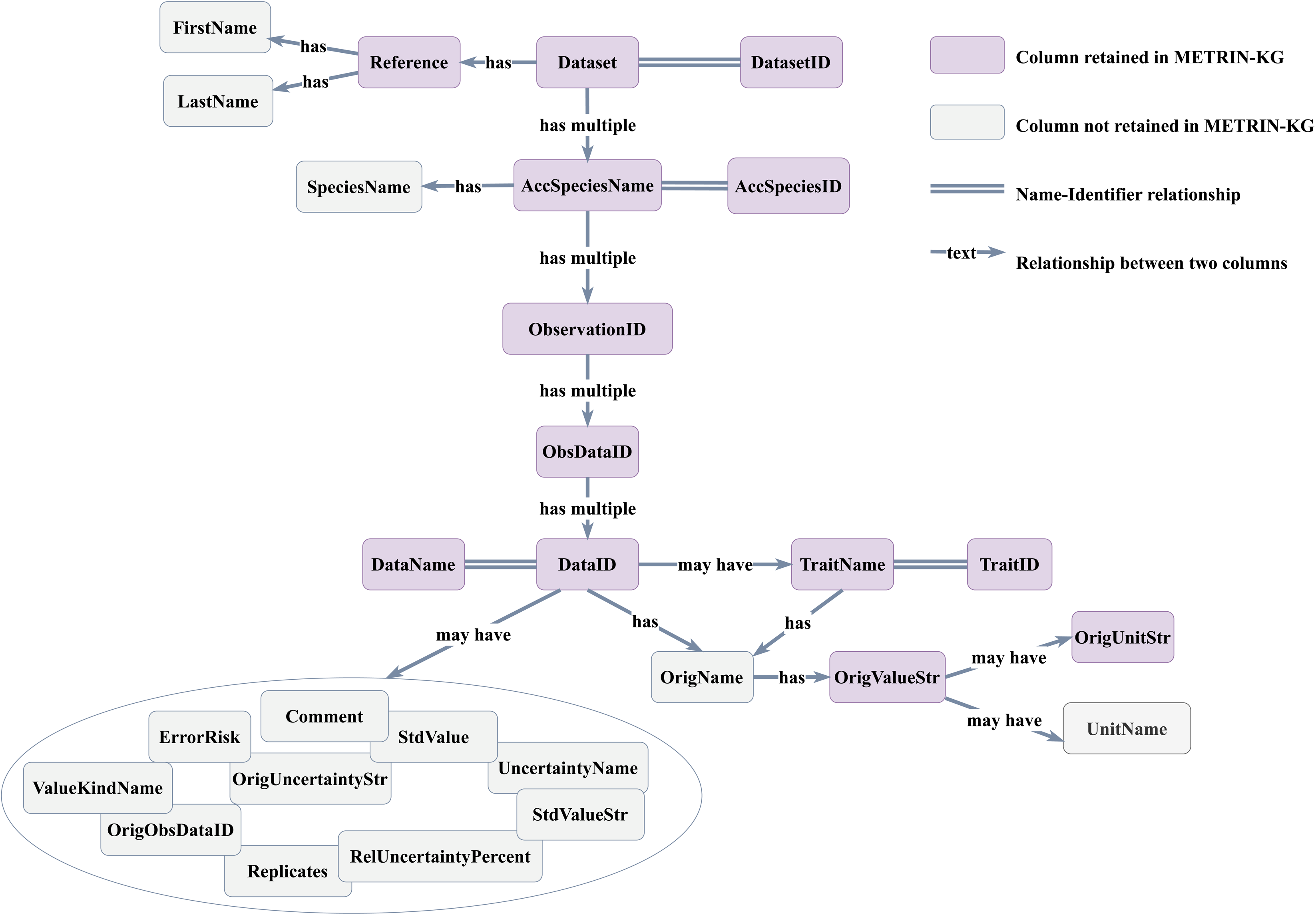
Original structure of traits dataset retrieved from TRY database. Purple-shaded rectangles represent columns retained for METRIN-KG. Gray-shaded rectangles represent columns not retained for METRIN-KG. Column relationships are represented by directional arrows.

The dataset was refined to a minimal form by removing columns that contained redundant or ambiguous information. For example, author first and last names were excluded, as this information is already available in the retained ‘Reference’ field. Likewise, the ‘OriginalName’ column duplicated data present in ‘DataName’ and was therefore omitted. ‘Replicates’ is also an entity mentioned in ‘DataName’ and therefore not retained. The TRY database includes standardized values for certain traits in the columns ‘ErrorRisk’, ‘StdValue’, ‘StdValueStr’, ‘OrigUncertainityStr’, ‘UncertaintyName’, ‘ValueKindName’, and ‘RelUncertainityPercent’. However, these were excluded due to their limited coverage across the dataset. This selective retention helped minimize redundancy and ensured consistency within the constructed knowledge graph. The metrics of the dataset following this refinement are as follows:

A) 65,675 unique species names (‘AccSpeciesName’; see **Supplementary Table 4**) out of 70,748 from TRY database were mapped to Wikidata (see section Taxonomy mapping).
B) Overall 1,826,445 traits & 17,212,303 non-trait (e.g.: number of replicates, latitude, longitude) records, data values, and corresponding units (**Supplementary Table 4**) were retained after mapping 65,672 species to Wikidata identifiers.

Query to retrieve metrics is described in [137].

### Mapping of GloBI data

The original structure and column descriptions of the GloBI data are described in **Figure-4** and **Supplementary Table 5**. The column descriptions are also provided on the github account of GloBI as depicted on its website [138]. Columns retained in the knowledge graph are indicated in the table and the figure. Lineage information was not retained, as it can be retrieved via federated queries through Wikidata. Physiological stage data was omitted due to the absence of reliable mappings to established ontologies or controlled vocabularies. The ‘eventDateTime’ field was removed owing to ambiguity - specifically, it was not clear whether it referred to the timing of the interaction event or the time of observation.

**Figure-4:**
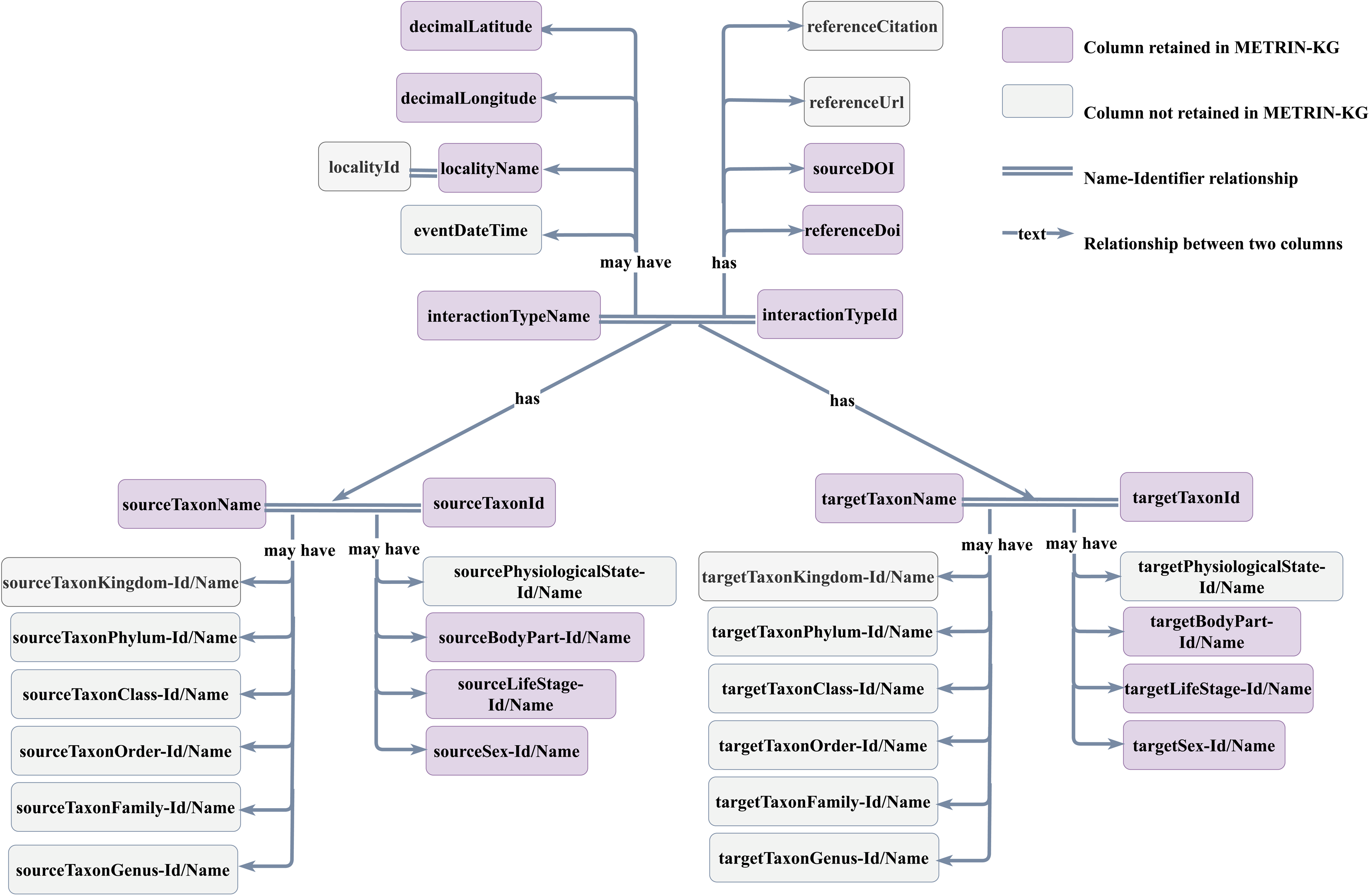
Original structure of interactions dataset retrieved from GloBI. Purple-shaded rectangles represent columns retained for METRIN-KG. Gray-shaded rectangles represent columns not retained for METRIN-KG. Column relationships are represented by directional arrows.

Overall, 20,480,925 records were obtained from 1,747,254 unique taxonomic identifiers and/or names (**Supplementary Table 4**), out of which 609,087 were mapped to 337,293 unique Wikidata identifiers. Overall, 12,872,681 GloBI records for only the mapped taxonomic identifiers were retained.

Query for the number of unique Wikidata identifiers and the number of records are provided in [139] and [140], respectively.

At the time of writing this manuscript, 1867 body part names were mapped to 996 ontology terms. 621 life stage names were mapped to 227 ontology terms. 57 biological sex names were mapped to 10 ontology terms.

### Bias

Biases in the mapped GloBI data in our knowledge graph reflect those documented for GloBI, including taxonomic and geographic skews, which are extensively discussed in the GloBI literature [76].

Metabolite coverage is primarily derived from two sources: (i) untargeted metabolomics data from approximately 1,600 plant extracts from the Pierre Fabre collection, as described in the ENPKG paper [69], and (ii) compound–taxon associations curated via Wikidata/LOTUS, which are also known to be biased toward well-studied taxa and compounds [68].

### Data re-use and case studies

#### Potential re-use and expansion of data

The current state of the knowledge graph provides a solid foundation. Its utility can be significantly enhanced by incorporating additional data sources and refining its structure.

##### a) Data re-use for studies in life sciences research

Researchers can query METRIN-KG to identify potential relationships between plant traits, interactions, and the presence or bioactivity of specific natural products. For example, one could query for plants with specific traits known to be associated with defense mechanisms and then explore the natural products they produce. The integrated data can be used to build predictive models for natural product occurrence or bioactivity based on plant traits and ecological context. This information can help prioritize plant species or natural product classes with a higher likelihood of possessing desired bioactivities. In addition, by analyzing trait-interaction-metabolome associations, researchers can prioritize plant species or ecological contexts that are more likely to yield novel or interesting natural products. Investigating such ecological roles of natural products can provide clues about their potential mechanisms of action and possible drug targets.

METRIN-KG can also facilitate the investigation of how natural products mediate species interactions. For instance, one could explore how specific compounds are associated with pollination or defense against herbivores in plants with particular traits. The understanding of the evolutionary pressures that shape the diversity of natural products can be investigated by linking them with the respective host traits and interactions.

In the context of potential data re-use for studies in life sciences research, we aimed to retrieve data from METRIN-KG on a few known questions in plant ecology, agriculture, and biodiversity. **Table-2** lists the questions and the respective links to the SPARQL queries.

##### b) User-friendly approach to add queries as examples within METRIN-KG

We devised a method for the users of METRIN-KG to propose a query and incorporate it within the examples (e.g. queries listed in **Table-2**) using the SPARQL query-editor proposed in [165,166]. Once a user proposes a query on EMI’s sparql-examples GitHub repository’s issues section, it will allow us to review it for inclusion in the examples. Once the review and corrections are complete, the query will be available on the SPARQL query endpoint [130]. While the examples listed in **Table-2** provide several complex queries, this option allows the users to provide context-specific queries useful for building research questions.

A tutorial for contributing queries to METRIN-KG is provided on its GitHub repository wiki [167].

##### c) Adding data or federating over knowledge graphs presenting datasets from other publicly available resources

Incorporating environmental data (e.g., climate data and soil data) could allow for the analysis of how environmental factors influence traits, interactions, and natural product production and distribution. Including data on the timing of biological events (e.g., flowering times) could add a temporal dimension to the analysis of interactions and natural product occurrence. In addition, incorporating image data (e.g., plant morphology) and spectral data (e.g., metabolomic profiles) could provide richer characterizations of the entities within the knowledge graph. Integrating broader ecological context, such as community composition and ecosystem dynamics, could provide a more holistic understanding of the relationships. Where ethically and legally appropriate, incorporating curated traditional knowledge related to plant uses and natural products could add valuable perspectives. Moreover, including data from other natural product databases (e.g.: PubChem [70], ChEMBL [71,168]) could broaden the coverage of chemical compounds and their properties.

##### d) Enhancing ontological structure and semantics

For developing more granular relationship types, moving beyond simple pairwise interactions to include more specific relationship types (e.g., pollination, herbivory, symbiosis) with associated properties (e.g., strength, specificity) could be useful. The first efforts in this direction are taken by GloBI [76] and BiotXplorer [77]. However, they are limited to pairwise interactions and not connected to other types of data, which our knowledge graph provides. A useful step in this direction could be to implement semantic reasoning, hence inferring new relationships and knowledge that are not explicitly stated in the data.

##### e) Democratizing METRIN-KG access: querying in natural language

In order to make available the knowledge in the METRIN-KG for non-SPARQL savvy users, we applied ExpasyGPT [169], a Large Language Model (LLM)-driven tool based on lightweight metadata that allows for querying knowledge graphs in natural language. This tool facilitates querying METRIN-KG as well as various other life-science databases in plain English. In other words, ExpasyGPT aids in constructing SPARQL queries in response to user’s text-format questions. Therefore, the final response is not given by the LLM but by querying METRIN-KG, which contains curated and high quality data, mitigating well-known LLM issues such as hallucinations, lack of domain specific knowledge and black-box behaviour. Consequently, this makes answers verifiable and reproducible. More precisely, ExpasyGPT implements a context-optimization approach that relies on two main sources of metadata: (i) pairs of question-query examples and (ii) automatically generated description of used classes and predicates. Both (i) and (ii) should be defined using well-known W3C vocabularies such as Vocabulary of Interlinked Datasets (VoID) and Shapes Constraint Language (SHACL), accessible through the SPARQL endpoint. Applying ExpasyGPT is straightforward thanks to the fact that the metadata already exists as described in **Table 2** and is automatically done. A tutorial of how to use ExpasyGPT is provided on METRIN-KG GitHub repository <u>wiki</u> [170].

**Table-2:**
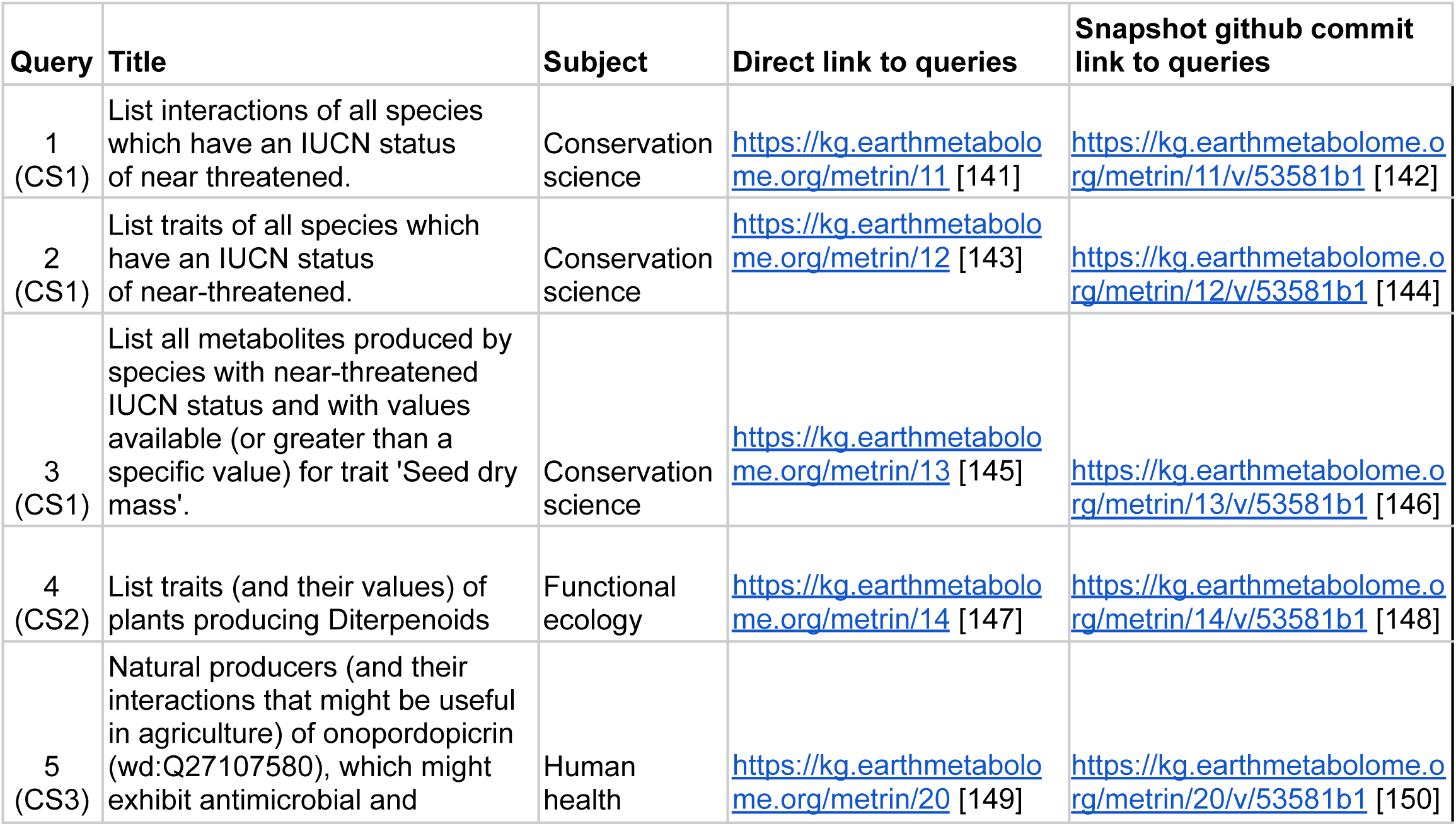
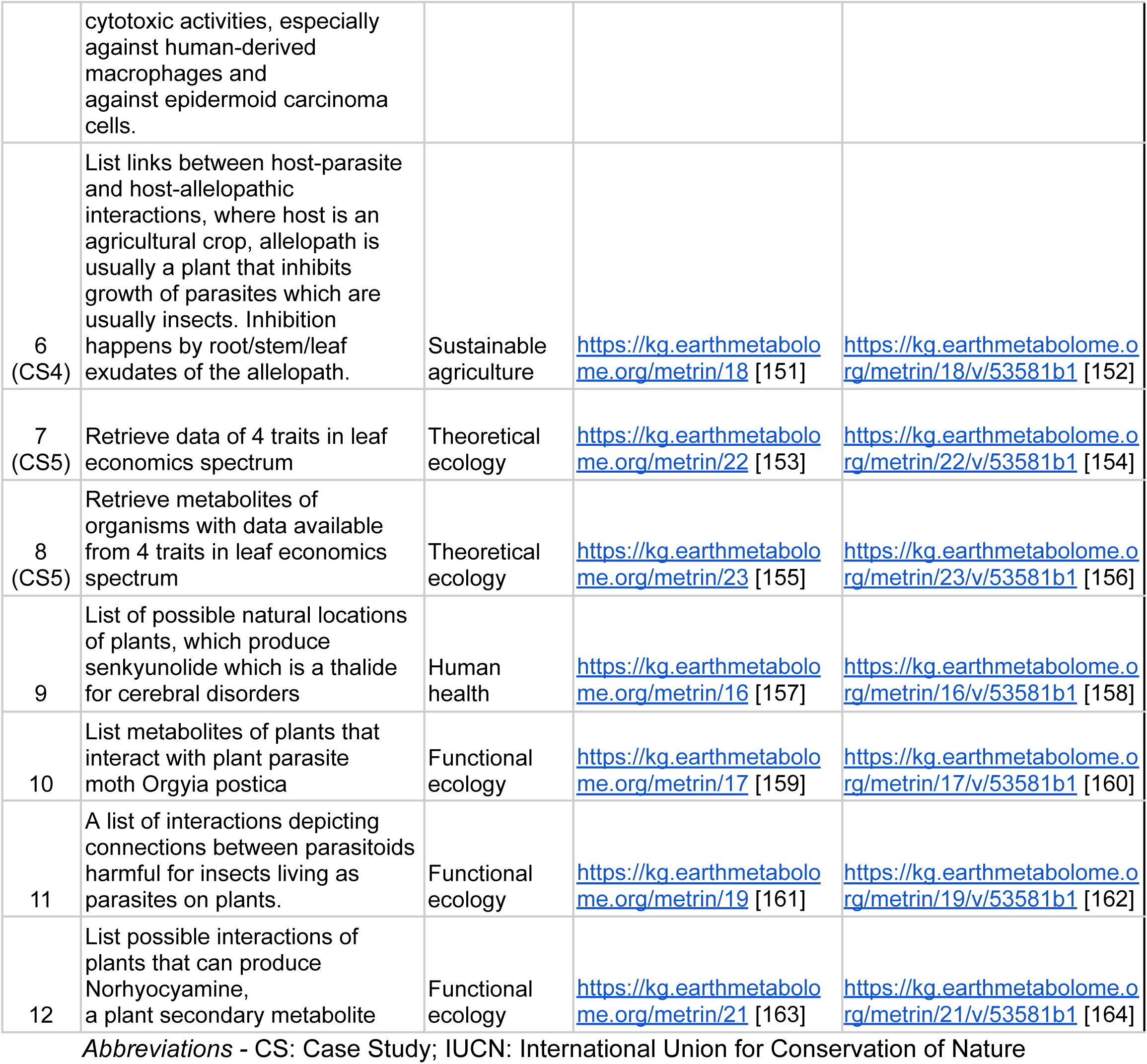
Examples of SPARQL queries for METRIN-KG and links to queries.

### Case studies

Out of the 12 examples listed in **Table-2**, we built 5 case study (CS) summaries for 8 as listed below.

#### CS1: List traits, interactions, and metabolites of all species which have an International Union for Conservation of Nature (IUCN) status of near threatened

Subject - Conservation science

The queries are described in [141], [143], [145] (**Table 2**)

Some studies have addressed the need to identify the threat status of plants by studying their functional traits [171–173]. Moreover, one study has suggested approaches for conservation of threatened plant species using functional trait patterns [174]. Some other studies have used predictive modelling to enhance knowledge of interacting species pairs (e.g.: predator-prey interactions) [175,176] in the context of their habitat and traits. In addition, some researchers have suggested and reviewed ecological restoration strategies for threatened ecosystems based on functional traits and plant-animal interactions [177–179].

Keeping such broad studies in mind, we developed 3 SPARQL queries retrieving the traits, interactions, and metabolites of plant species with IUCN status of near threatened. Such queries would aid the researchers in collecting data relevant to near threatened plant species (in some cases, insects) and develop strategies to model patterns specific to climate change, thereby developing ecological restoration plans.

Basic metrics obtained from the results of the queries are provided in **Table 3**. The results obtained from the three queries contain 9,299 unique species with highly heterogeneous data coverage. While interaction data were available for the majority of species (8,598), trait measurements were substantially sparser (1,051 species, 19,279 measurements across 98 traits), and metabolite data were limited to only 47 species (818 unique InChIKeys and 1,000 records). 37 species possessed all three data types.

**Table-3:** Metrics for CS1.

| Metric | Value |
| --- | --- |
| Total unique species (all data) | 9299 |
| Unique species with trait data | 1051 |
| Unique species with metabolite data | 47 |
| Unique species with interaction data | 8598 |
| Unique species with traits and metabolites | 47 |
| Unique species with all three data types | 37 |
| Total unique traits | 98 |
| Total unique metabolites (InChIKeys) | 818 |
| Total unique interaction types | 24 |
| Total trait measurements | 19279 |
| Total metabolite records | 1000 |
| Total interaction records | 12396 |

While these 3 queries can be combined into one, we kept them separate to improve efficiency and retain modularity.

#### CS2: List traits and their values for plants producing diterpenoids

Subject - Functional ecology

The query is described in [147] (**Table 2**).

Recent studies have utilized trait data to understand the implications of diterpenoids producing invasive plant species and their allelopathic relationships with agricultural crops [180–185]. In the context of agriculture, there have been reviews of plant traits and their effects on insect herbivory [186], crop resilience (e.g.: rice) of plants producing diterpenoids [187–189], and the implications of drought stress on diterpenoid producing plants and their traits [190]. In the context of biodiversity conservation, there have been studies on plant traits and their metabolic profile [191]. In addition, there has been research on plant interactions with microbial communities of plants where leaf surface has been reported to contain diterpenoids (tobacco and phyllosphere) [192].

Considering such diverse research on diterpenoids-producing plants or where diterpenoids are known to be extracted from plant surfaces, we developed a simple query to list traits of plants which have been known to produce or harbor diterpenoids.

**Figure-5** provides the trait coverage of top 30 species selected based on measurement frequency from the results. **Supplementary Tables 6** and **7** provide complete summaries of species and trait distributions. The dataset contains 98 traits measured across 2,131 species, with photosynthesis per leaf area being the most frequently measured trait (16,472 measurements) followed by leaf nitrogen content (14,841 measurements) and specific leaf area (14,698 measurements). *Pinus sylvestris* has the highest number of trait measurements (14,557 records), followed by *Qualea grandiflora* (5,260 records) and *Fagus sylvatica* (4,978 records).

**Figure-5:**
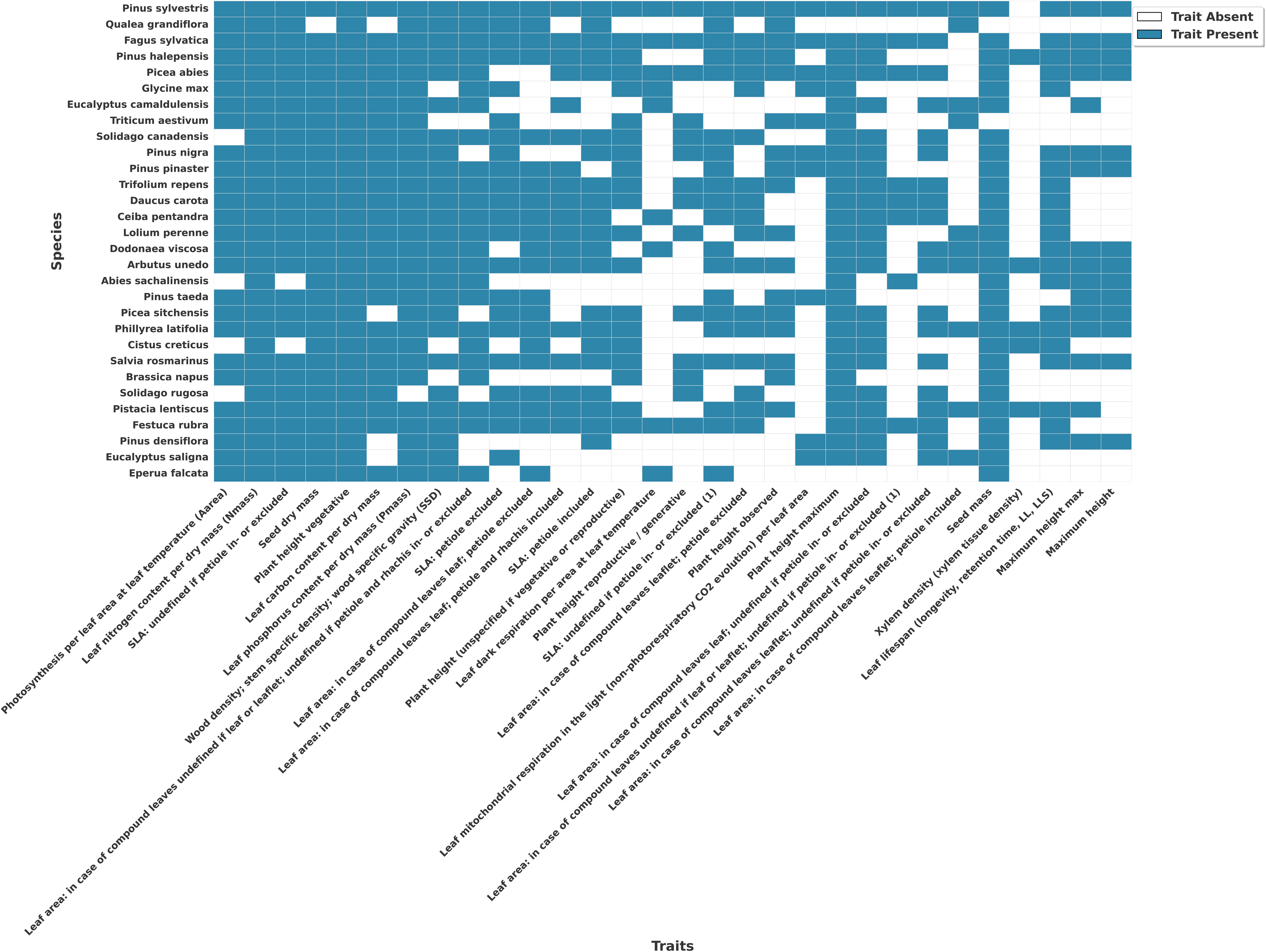
CS2 - Trait coverage of top 30 species selected based on measurement frequency from the results. Blue shows measured traits for a species; white shows no trait measured.

#### CS3: Natural producers and their biotic interactions that might be useful in agriculture of onopordopicrin

Subject - Human health

The query is described in [149] (**Table 2**).

Onopordopicrin is a metabolite produced by several plants like *Arctium lappa* and genus *Shangwua.* It has been shown to potentially exhibit antimicrobial and cytotoxic activities, especially against human-derived macrophages and against epidermoid carcinoma cells [193]. There is less scientific evidence to support these claims and research in this area has only recently been enhanced [194–197], making onopordopicrin an appropriate and timely subject for prospective research studies in human health and pharmacology.

To specifically target the research in this area, we developed a query to retrieve the natural producers of onopordopicrin and their biotic interactions.

The results comprise of 98 diterpenoid-producing species connected through 101 biotic interactions. While we did not find *Arctium lappa* and genus *Shangwua* in the results, *Eriophyllum confertiflorum*, emerges as a hub species with 77 incoming connections, that interacts with numerous other organisms (**Figure-6**). The network shows a highly skewed degree distribution where a small number of plant species (like *Centaurea melitensis* with 14 connections) dominate interactions, while the majority of species participate in only one or two interactions each.

**Figure-6:**
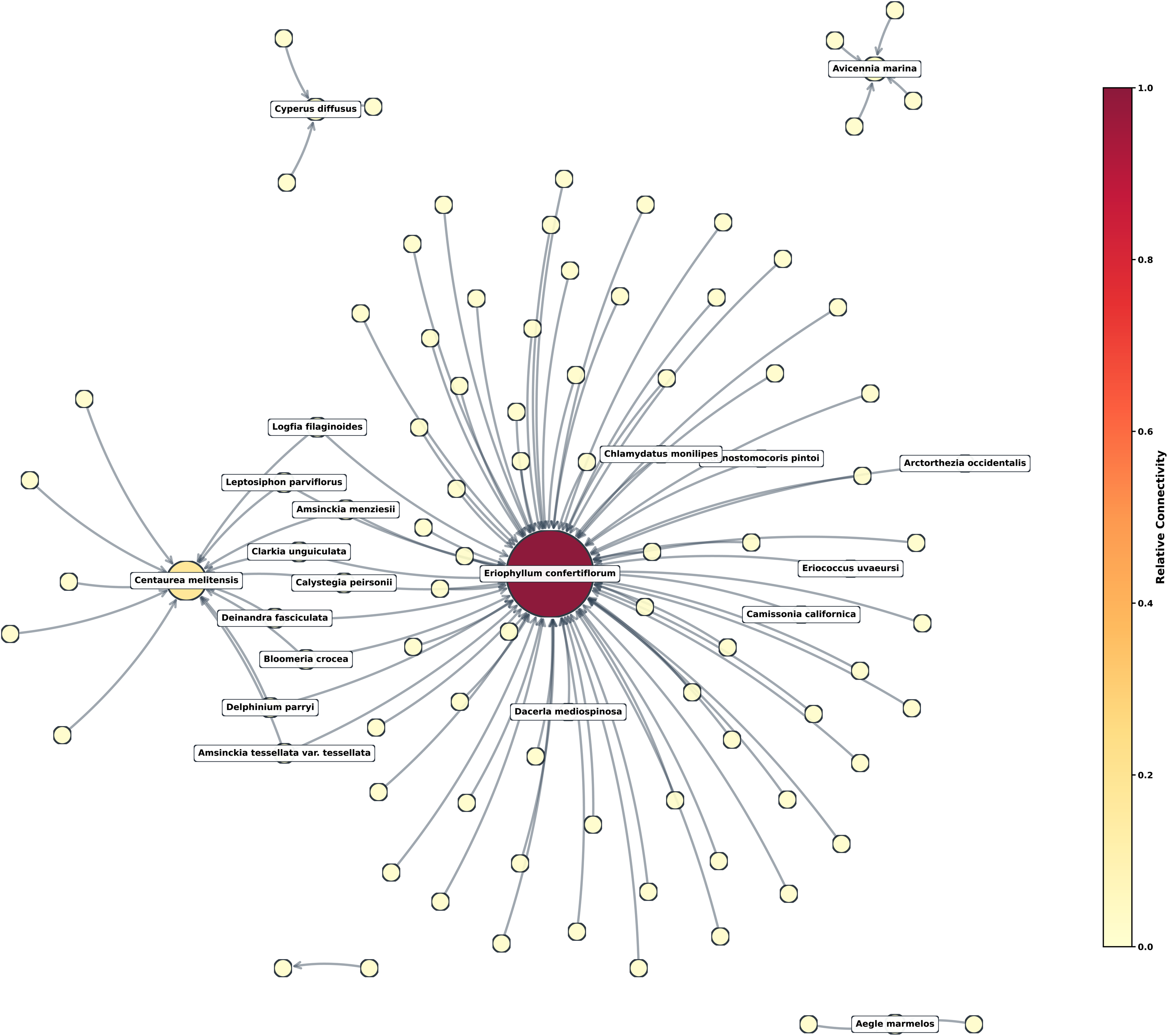
CS3 - Interaction network for diterpenoid producing plant species. Node size and color represent connection count. Only top-20 species with the highest connection count are labelled.

#### CS4: Allelopathy interactions in push-pull agriculture

Subject - Sustainable agriculture

The query is described in [151] (**Table 2**).

Studies on allelopathy have advanced our knowledge of plant interactions with other organisms, particularly in agriculture [198–200] and invasion ecology [180]. For instance, species of genera *Desmodium*, through their root and stem exudates, may protect crops of maize (*Zea mays*) and sorghum (*Sorghum*) against attack from stemborer arthropods and weed *Striga hermonthica* [201,202]. Besides focussed research [203], there are few studies exploring the allelopathic properties and the plant metabolites responsible for them.

We developed a query to target such interactions and the metabolites produced by the protective allelopathic plants.

In the original GloBI dataset, several fungal genera (e.g., *Fusarium*, *Puccinia*, *Claviceps, Sclerotinia sclerotiorum*) were misidentified as allelopath, likely resulting from automated inverse relationship generation where Plant ‘hasPathogen’ Fungus was inverted to Fungus ‘allelopathOf’ Plant. In the result obtained for CS4, we removed 55 interactions (45% of the original dataset) where fungi were incorrectly classified as allelopaths, as these organisms represent direct pathogenic relationships rather than plant-mediated chemical interference. After applying this filter, the results show 67 tripartite interactions involving 46 parasites, 11 allelopathic plants, and 10 crops (**Figure-7, Supplementary Table 8**).

**Figure-7:**
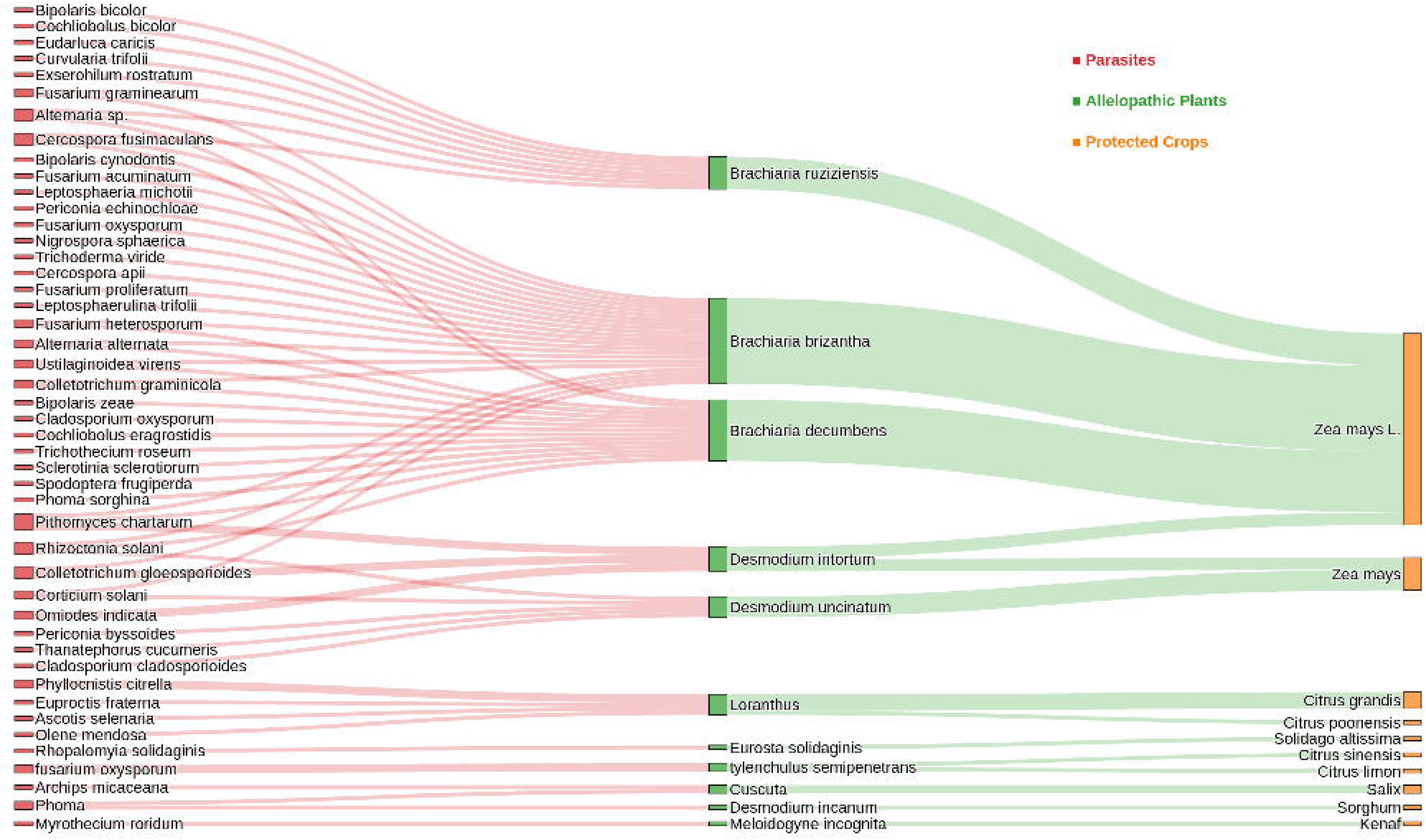
CS4 - Sankey plot representing the flow of allelopathy interactions in push-pull agriculture. Parasites are shown in red, allelopathic plants in green, and protected crops in orange.

#### CS5: Retrieve traits and metabolites for organisms for which traits from Leaf Economic Spectrum (LES) are available

Subject - Theoretical ecology

Queries are described in [153] and [155] (**Table 2**).

Leaf Economic Spectrum is a foundational concept in plant ecology, describing coordinated variation in leaf traits across species, typically forming a trade-off between resource-acquisitive and resource-conservative strategies [81,204]. Resource-acquisitive species have traits like high specific leaf area (SLA), high nitrogen content, short leaf lifespan, which are optimized for rapid growth in resource-rich environments. Resource-conservative species show traits like low SLA, low nutrient content, long leaf lifespan, which are better suited for resource-poor environments with conservative growth. These relationships are global, cross-taxonomic, and continuous. They help predict ecosystem functions, species distributions, and responses to environmental changes.

We developed a query to retrieve values for four traits (including their sub-categories totalling to six) relevant for LES - specific leaf area, leaf dry matter content, leaf nitrogen content, and lead phosphorus content. Further, we developed another query to retrieve metabolites produced by species with data for four LES traits available in the knowledge graph. While we were unable to identify plant species with both metabolite data and complete trait coverage across all four categories, this limitation itself reinforces the value of our knowledge graph approach and the importance of continued data integration efforts.

**Table-4** depicts LES trait data from 1,058 plant species with metabolite profiles from 627 species, identifying 49 species (with incomplete trait coverage) with both datasets available for comparative analysis. The data encompassed 6 key functional traits across 2,324 measurements and 11,313 unique metabolites across 16,587 associations.

**Table-4:**
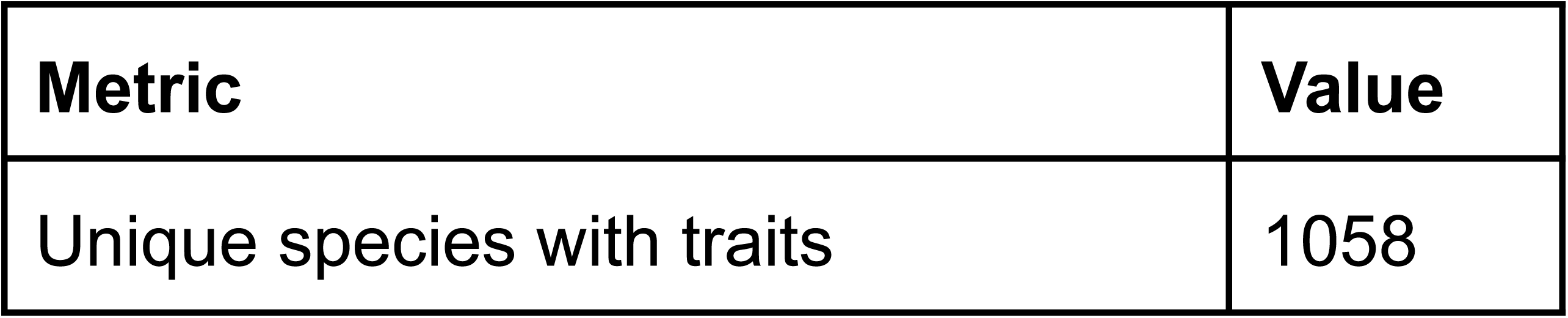
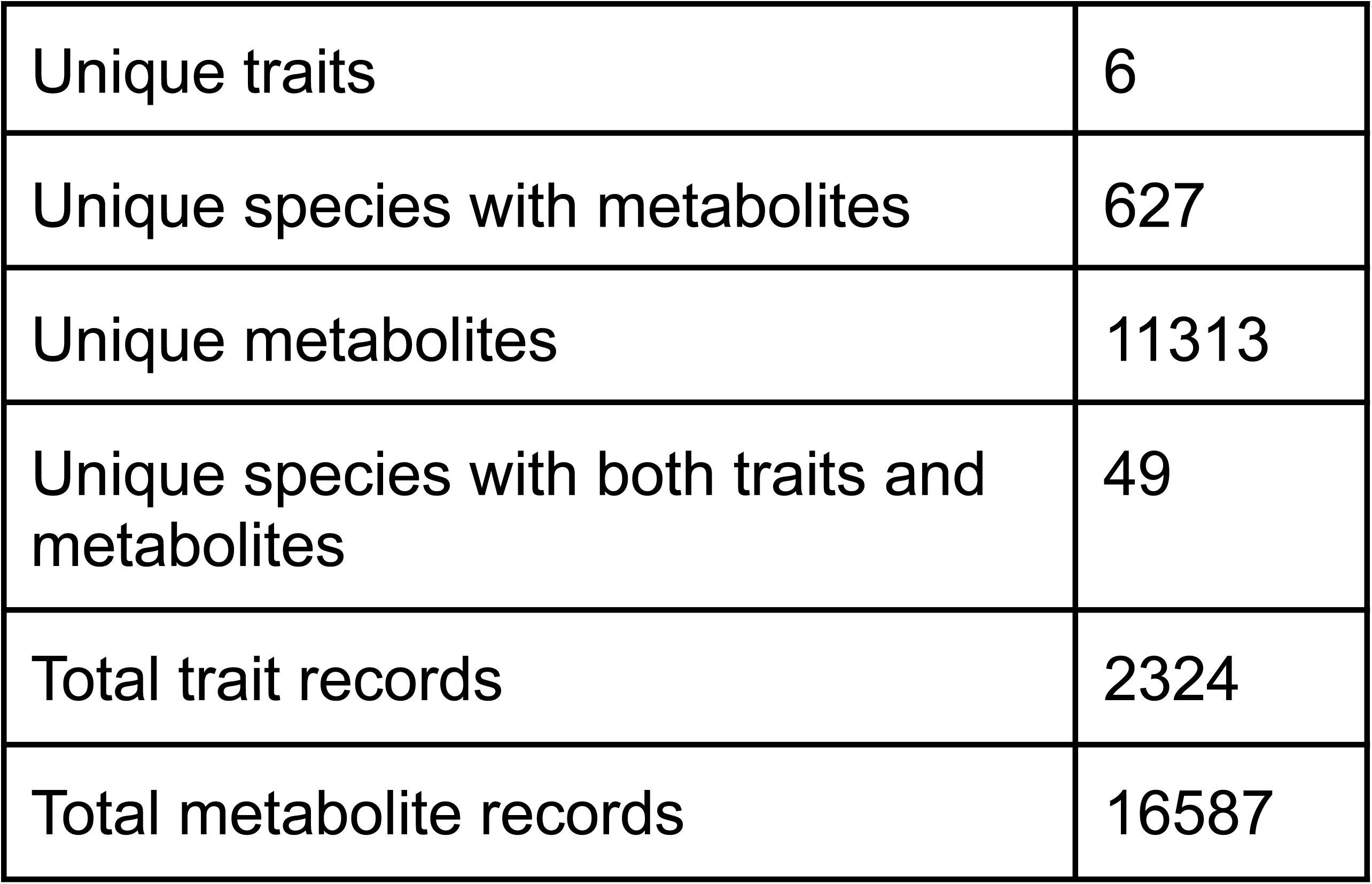
Metrics for CS4.

At the time of writing this manuscript, the results of our queries reflect the state of METRIN-KG and the federated sources available. As these underlying datasets are continuously updated and expanded, the query outputs presented here may evolve over time. All scripts used to generate the tables and figures for case studies are available at METRIN-KG GitHub repository [128].

## Conclusions

We provide METRIN-KG aiming to integrate seemingly disconnected databases of plant metabolomes, plant traits, and interactions. To facilitate exploration of this resource, we provide a way to search for those connections through a user interface. We aspire to draw the attention of researchers in areas of drug-discovery, ecology, human health, biodiversity conservation, and agriculture to the power of knowledge graphs in integrating life-sciences datasets. We wish to increase awareness amongst researchers to include as varied data as possible, so as to approach research questions from multiple perspectives. In this context, we provide representative case studies to retrieve data for different research projects. We plan to extend METRIN-KG to incorporate the full breadth of metabolome data from plants in the near future. In the distant long-term, we plan to extend it, adding more information on metabolite data for other kingdoms beyond Plantae, while updating the trait and interaction data from TRY and GloBI, respectively, as new versions are released. We hope that our efforts will encourage other researchers to use and contribute to this resource.

## Availability of source code and requirements

a) METRIN-KG Project name: metrin-kg Project homepage: https://github.com/earth-metabolome-initiative/metrin-kg License: GPL-3.0 license Operating system(s): Ubuntu 24.04.2 LTS Programming language: Python Hardware requirements: 32 GB RAM, 500 GB disk-capacity RRID: SCR_027914 biotools: metrin_kg METRIN-KG’s code is also available through Zenodo [205].
b) Earth Metabolome Initiative Ontology Project name: Earth Metabolome Initiative Ontology, illustrated with an example of knowledge-graph construction Project homepage: https://github.com/earth-metabolome-initiative/earth_metabolome_ontology License: Software code under GPL-3.0 license, Ontology under CC0-1.0 license Operating system(s): modern Linux/Windows/Unix distribution (64-bit OS) Programming language: OWL, Python Hardware requirements: any x86_64 or ARM64 CPU with at least 2 cores, and a minimum of 8 GB RAM and 60 GB disk are recommended for functional setups RRID: SCR_027915 biotools: earth_metabolome_initiative_ontology

## Data availability

All input and output files used in mapping taxonomy, metadata, ontology, and knowledge graph construction as well as RDF files are available on EMI’s Zenodo repository for METRIN-KG (CC0-1.0) [104] and TRY (CC-BY-4.0) [95].

## Abbreviations

AO: Agronomy Ontology
APO: Ascomycete Phenotype Ontology
BRENDA: Braunschweig Enzyme Database
BSPO: Biological Spatial Ontology
BTO: BRENDA Tissue Ontology
CEPH: Cephalopod Ontology
ChEBI: Chemical Entities of Biological Interest
CL: Cell Ontology
CLAO: Collembola Anatomy Ontology
CS: Case Study
CSV: Comma-Separated Values
DBGI: Digital Botanical Gardens Initiative
DDANAT: Dictyostelium Discoideum Anatomy
ECOCORE: Ecology Core Ontology
EFO: Experimental Factor Ontology
EMI: Earth Metabolome Initiative
ENPKG: Experimental Natural Products Knowledge Graph
ENVO: Environment Ontology
FAIR: Findable, Accessible, Interoperable, and Reusable
FAO: Fungal Anatomy Ontology
FLOPO: Flora Phenotype Ontology
FMA: Foundational Model of Anatomy
FOODON: Food Ontology
GallOnt: ontology for plant gall phenotypes
GloBI: Global Biotic Interactions
GO: Gene Ontology
HAO: Hymenoptera Anatomy Ontology
IDO: Infectious Disease Ontology
IDOMAL: Infectious Disease Ontology for Malaria
IRI: Internationalized Resource Identifier
IUCN: International Union for Conservation of Nature
KG: Knowledge Graph
LES: Leaf Economic Spectrum
LMA: Leaf dry mass per area
LOTUS: *not an acronym*
METRIN-KG: MEtabolomes, TRaits, and INteractions-Knowledge Graph
NCIT: National Cancer Institute Thesaurus
OBI: Ontology for Biomedical Investigations
OGC: Open Geospatial Consortium
OMIT: Ontology for MIcroRNA Target Prediction
OWL: Web Ontology Language
PATO: Phenotype and Trait Ontology
PHIPO: Pathogen-Host Interaction Phenotype Ontology
PNUE: Photosynthetic Nitrogen Use Efficiency
PO: Plant Ontology
PORO: Porifera Ontology
QUDT: Quantities, Units, Dimensions, and Types
RDF: Resource Description Framework
RO: Relation Ontology
SHACL: Shapes Constraint Language
SKOS: Simple Knowledge Organization System
SLA: Specific Leaf Area
SOSA: Sensor, Observation, Sample, and Actuator
SPARQL: SPARQL Protocol and RDF Query Language
SSD: Stem Specific Density
SQL: Structured Query Language
TGMA: Mosquito gross anatomy ontology
TPU: Triose Phosphate Utilisation
TRY: *not an acronym*
TSV: Tab-Separated Values
UBERON: Uber-Anatomy Ontology
UI: User Interface
URL: Uniform Resource Locators
VoID: Vocabulary of Interlinked Datasets
W3C: World Wide Web Consortium
WGS84: World Geodetic System 1984
WUE: Water Use Efficiency

## Authors’ Contributions

D.T., P.M.A., T.M.F., and E.D. conceptualized the study. D.T. and T.M.F curated the data, designed the methodology and developed the knowledge graph. T.M.F. designed and developed the Earth Metabolome Initiative Ontology with inputs from D.T. and P.M.A.. D.T. and T.M.F. prepared the visualization. D.T. and E.D. identified the example case-studies relevant to the knowledge graph. T.M.F and P.M.A. provided computing resources for the project. P.M.A, T.M.F., and E.D. acquired the funding for the project. D.T. and T.M.F. wrote the original draft. All authors reviewed and edited the manuscript.

## Acknowledgements

The authors thank members of Earth Metabolome Initiative consortium for their useful comments and suggestions on the project.

## Supplementary Table titles and legends

**Supplementary Table 1: List of taxonomy databases used in GloBI**

**Supplementary Table 2: List of ontologies used in METRIN-KG.**

**Supplementary Table 3**: **Column descriptions of data retrieved from TRY database.** Column names and descriptions were retained from the files provided by TRY database managers

**Supplementary Table 4: Metrics for TRY database and GloBI.**

**Supplementary Table 5**: **Column descriptions of data retrieved from GloBI.** Column names and descriptions obtained from GloBI dataset template repository [158].

**Supplementary Table 6: CS2 - Summary of species distributions.**

**Supplementary Table 7: CS2 - Summary of trait distributions.**

**Supplementary Table 8: Filtered data without fungi species for CS4.**

## References

1. Hortal J, Bello F de, Diniz-Filho JAF, Lewinsohn TM, Lobo JM, Ladle RJ. Seven Shortfalls that Beset Large-Scale Knowledge of Biodiversity. Annu Rev Ecol Evol Syst. Annual Reviews; 2015; doi: 10.1146/annurev-ecolsys-112414-054400.

2. Cardoso P, Erwin T, Borges P, New T. The seven impediments in invertebrate conservation and how to overcome them. Biol Conserv. 2011; doi: 10.1016/j.biocon.2011.07.024.

3. Pollock LJ, Kitzes J, Beery S, Gaynor KM, Jarzyna MA, Mac Aodha O, et al.. Harnessing artificial intelligence to fill global shortfalls in biodiversity knowledge. Nat Rev Biodivers. Nature Publishing Group; 2025; doi: 10.1038/s44358-025-00022-3.

4. Walker TWN, Alexander JM, Allard P-M, Baines O, Baldy V, Bardgett RD, et al.. Functional Traits 2.0: The power of the metabolome for ecology. J Ecol. 2022; doi: 10.1111/1365-2745.13826.

5. Hilker M. New Synthesis: Parallels Between Biodiversity and Chemodiversity. J Chem Ecol. 2014; doi: 10.1007/s10886-014-0402-8.

6. Schuman MC, Baldwin IT. The Layers of Plant Responses to Insect Herbivores. Annu Rev Entomol. Annual Reviews; 2016; doi: 10.1146/annurev-ento-010715-023851.

7. Wink M. Evolution of secondary metabolites from an ecological and molecular phylogenetic perspective. Phytochemistry. 2003; doi: 10.1016/S0031-9422(03)00300-5.

8. Hart CE, Gadiya Y, Kind T, Krettler CA, Gaetz M, Misra BB, et al.. Defining the limits of plant chemical space: challenges and estimations. GigaScience. 2025; doi: 10.1093/gigascience/giaf033.

9. Van Dam NM, Van Der Meijden E. A Role for Metabolomics in Plant Ecology. In: Hall RD, editor. Annu Plant Rev Vol 43. 1st ed. Wiley; 2011; doi: 10.1002/9781444339956.ch4.

10. Weckwerth W. Metabolomics in Systems Biology. Annu Rev Plant Biol. Annual Reviews; 2003; doi: 10.1146/annurev.arplant.54.031902.135014.

11. Echeverri A, Karp DS, Naidoo R, Tobias JA, Zhao J, Chan KMA. Can avian functional traits predict cultural ecosystem services? People Nat. 2020; doi: 10.1002/pan3.10058.

12. Wong MKL, Guénard B, Lewis OT. Trait-based ecology of terrestrial arthropods. Biol Rev. 2019; doi: 10.1111/brv.12488.

13. Lundgren EJ, Schowanek SD, Rowan J, Middleton O, Pedersen RØ, Wallach AD, et al.. Functional traits of the world’s late Quaternary large-bodied avian and mammalian herbivores. Sci Data. Nature Publishing Group; 2021; doi: 10.1038/s41597-020-00788-5.

14. Kunin WE, Vergeer P, Kenta T, Davey MP, Burke T, Ian Woodward F, et al.. Variation at range margins across multiple spatial scales: environmental temperature, population genetics and metabolomic phenotype. Proc R Soc B Biol Sci. 2009; doi: 10.1098/rspb.2008.1767.

15. Dwyer JM, Hobbs RJ, Mayfield MM. Specific leaf area responses to environmental gradients through space and time. Ecology. 2014; doi: 10.1890/13-0412.1.

16. Adler PB, Salguero-Gómez R, Compagnoni A, Hsu JS, Ray-Mukherjee J, Mbeau-Ache C, et al.. Functional traits explain variation in plant life history strategies. Proc Natl Acad Sci. Proceedings of the National Academy of Sciences; 2014; doi: 10.1073/pnas.1315179111.

17. Pistón N, de Bello F, Dias ATC, Götzenberger L, Rosado BHP, de Mattos EA, et al.. Multidimensional ecological analyses demonstrate how interactions between functional traits shape fitness and life history strategies. J Ecol. 2019; doi: 10.1111/1365-2745.13190.

18. Walker TWN, Weckwerth W, Bragazza L, Fragner L, Forde BG, Ostle NJ, et al.. Plastic and genetic responses of a common sedge to warming have contrasting effects on carbon cycle processes. Ecol Lett. 2019; doi: 10.1111/ele.13178.

19. van der Plas F, Schröder-Georgi T, Weigelt A, Barry K, Meyer S, Alzate A, et al.. Plant traits alone are poor predictors of ecosystem properties and long-term ecosystem functioning. Nat Ecol Evol. Nature Publishing Group; 2020; doi: 10.1038/s41559-020-01316-9.

20. Firn J, McGree JM, Harvey E, Flores-Moreno H, Schütz M, Buckley YM, et al.. Leaf nutrients, not specific leaf area, are consistent indicators of elevated nutrient inputs. Nat Ecol Evol. Nature Publishing Group; 2019; doi: 10.1038/s41559-018-0790-1.

21. Laughlin DC, Gremer JR, Adler PB, Mitchell RM, Moore MM. The Net Effect of Functional Traits on Fitness. Trends Ecol Evol. Elsevier; 2020; doi: 10.1016/j.tree.2020.07.010.

22. Eisenhauer N, Bonfante P, Buscot F, Cesarz S, Guerra C, Heintz-Buschart A, et al.. Biotic Interactions as Mediators of Context-Dependent Biodiversity-Ecosystem Functioning Relationships. Res Ideas Outcomes. Pensoft Publishers; 2022; doi: 10.3897/rio.8.e85873.

23. Maestre FT, Bowker MA, Escolar C, Puche MD, Soliveres S, Maltez-Mouro S, et al.. Do biotic interactions modulate ecosystem functioning along stress gradients? Insights from semi-arid plant and biological soil crust communities. Philos Trans R Soc B Biol Sci. 2010; doi: 10.1098/rstb.2010.0016.

24. Jayaramaiah RH, Egidi E, Macdonald CA, Singh BK. Linking biodiversity and biotic interactions to ecosystem functioning. J Sustain Agric Environ. 2024; doi: 10.1002/sae2.12119.

25. de Vries S, Feussner I. Biotic interactions, evolutionary forces and the pan-plant specialized metabolism. Philos Trans R Soc B Biol Sci. Royal Society; 2024; doi: 10.1098/rstb.2023.0362.

26. Defossez E, Pitteloud C, Descombes P, Glauser G, Allard P-M, Walker TWN, et al.. Spatial and evolutionary predictability of phytochemical diversity. Proc Natl Acad Sci. Proceedings of the National Academy of Sciences; 2021; doi: 10.1073/pnas.2013344118.

27. Agrawal AA, Fishbein M, Halitschke R, Hastings AP, Rabosky DL, Rasmann S. Evidence for adaptive radiation from a phylogenetic study of plant defenses. Proc Natl Acad Sci. Proceedings of the National Academy of Sciences; 2009; doi: 10.1073/pnas.0904862106.

28. Ehrlich PR, Raven PH. Butterflies and Plants: A Study in Coevolution. Evolution. [Society for the Study of Evolution, Wiley]; 1964; doi: 10.2307/2406212.

29. Bennett RN, Wallsgrove RM. Secondary metabolites in plant defence mechanisms. New Phytol. 1994; doi: 10.1111/j.1469-8137.1994.tb02968.x.

30. Walker TWN, Schrodt F, Allard P-M, Defossez E, Jassey VEJ, Schuman MC, et al.. Leaf metabolic traits reveal hidden dimensions of plant form and function. Sci Adv. American Association for the Advancement of Science; 2023; doi: 10.1126/sciadv.adi4029.

31. Deng P, Yin R, Wang H, Chen L, Cao X, Xu X. Comparative analyses of functional traits based on metabolome and economic traits variation of Bletilla striata: Contribution of intercropping. Front Plant Sci. Frontiers; 2023; doi: 10.3389/fpls.2023.1147076.

32. Wei J, Wang A, Li R, Qu H, Jia Z. Metabolome-wide association studies for agronomic traits of rice. Heredity. Nature Publishing Group; 2018; doi: 10.1038/s41437-017-0032-3.

33. Delory BM, Callaway RM, Semchenko M. A trait-based framework linking the soil metabolome to plant–soil feedbacks. New Phytol. 2024; doi: 10.1111/nph.19490.

34. Shi T, Zhu A, Jia J, Hu X, Chen J, Liu W, et al.. Metabolomics analysis and metabolite-agronomic trait associations using kernels of wheat (Triticum aestivum) recombinant inbred lines. Plant J. 2020; doi: 10.1111/tpj.14727.

35. Green SJ, Brookson CB, Hardy NA, Crowder LB. Trait-based approaches to global change ecology: moving from description to prediction. Proc R Soc B Biol Sci. Royal Society; 2022; doi: 10.1098/rspb.2022.0071.

36. Walter HE, Pagel J, Cooksley H, Neu A, Schleuning M, Schurr FM. Effects of biotic interactions on plant fecundity depend on spatial and functional structure of communities and time since disturbance. J Ecol. 2023; doi: 10.1111/1365-2745.14018.

37. Pueyo Y, Kéfi S, Díaz-Sierra R, Alados CL, Rietkerk M. The role of reproductive plant traits and biotic interactions in the dynamics of semi-arid plant communities. Theor Popul Biol. 2010; doi: 10.1016/j.tpb.2010.09.001.

38. Gaüzère P, O’Connor L, Botella C, Poggiato G, Münkemüller T, Pollock LJ, et al.. The diversity of biotic interactions complements functional and phylogenetic facets of biodiversity. Curr Biol. 2022; doi: 10.1016/j.cub.2022.03.009.

39. Tenenboim H, Brotman Y. Omic Relief for the Biotically Stressed: Metabolomics of Plant Biotic Interactions. Trends Plant Sci. 2016; doi: 10.1016/j.tplants.2016.04.009.

40. Gupta S, Schillaci M, Roessner U. Metabolomics as an emerging tool to study plant–microbe interactions. Emerg Top Life Sci. 2022; doi: 10.1042/ETLS20210262.

41. Henderson D, Sedio BE, Tello JS, Cayola L, Fuentes AF, Alvestegui B, et al.. Ecological metabolomics of tropical tree communities across an elevational gradient: Implications for chemically-mediated biotic interactions and species diversity. bioRxiv. 2023; doi: 10.1101/2023.10.04.560880.

42. Maag D, Erb M, Glauser G. Metabolomics in plant–herbivore interactions: challenges and applications. Entomol Exp Appl. 2015; doi: 10.1111/eea.12336.

43. Majumdar S, Kaur H, Rinella MJ, Kundu A, Vadassery J, Erbilgin N, et al.. Synergistic effects of canopy chemistry and autogenic soil biota on a global invader. J Ecol. 2023; doi: 10.1111/1365-2745.14113.

44. Semchenko M, Nettan S, Sepp A, Zhang Q, Abakumova M, Davison J, et al.. Soil biota and chemical interactions promote co-existence in co-evolved grassland communities. J Ecol. 2019; doi: 10.1111/1365-2745.13220.

45. Burghardt KT, Bradford MA, Schmitz OJ. Acceleration or deceleration of litter decomposition by herbivory depends on nutrient availability through intraspecific differences in induced plant resistance traits. J Ecol. 2018; doi: 10.1111/1365-2745.13002.

46. Heinen R, Biere A, Bezemer TM. Plant traits shape soil legacy effects on individual plant–insect interactions. Oikos. 2020; doi: 10.1111/oik.06812.

47. De Long JR, Heinen R, Hannula SE, Jongen R, Steinauer K, Bezemer TM. Plant-litter-soil feedbacks in common grass species are slightly negative and only marginally modified by litter exposed to insect herbivory. Plant Soil. 2023; doi: 10.1007/s11104-022-05590-3.

48. Eissenstat DM, Kucharski JM, Zadworny M, Adams TS, Koide RT. Linking root traits to nutrient foraging in arbuscular mycorrhizal trees in a temperate forest. New Phytol. 2015; doi: 10.1111/nph.13451.

49. Stiblíková P, Klimeš A, Cahill JF, Koubek T, Weiser M. Interspecific differences in root foraging precision cannot be directly inferred from species’ mycorrhizal status or fine root economics. Oikos. 2023; doi: 10.1111/oik.08995.

50. Xia M, Valverde-Barrantes OJ, Suseela V, Blackwood CB, Tharayil N. Coordination between compound-specific chemistry and morphology in plant roots aligns with ancestral mycorrhizal association in woody angiosperms. New Phytol. 2021; doi: 10.1111/nph.17561.

51. Kong C-H, Zhang S-Z, Li Y-H, Xia Z-C, Yang X-F, Meiners SJ, et al.. Plant neighbor detection and allelochemical response are driven by root-secreted signaling chemicals. Nat Commun. Nature Publishing Group; 2018; doi: 10.1038/s41467-018-06429-1.

52. Li L, Li S-M, Sun J-H, Zhou L-L, Bao X-G, Zhang H-G, et al.. Diversity enhances agricultural productivity via rhizosphere phosphorus facilitation on phosphorus-deficient soils. Proc Natl Acad Sci. Proceedings of the National Academy of Sciences; 2007; doi: 10.1073/pnas.0704591104.

53. Steinauer K, Thakur MP, Emilia Hannula S, Weinhold A, Uthe H, van Dam NM, et al.. Root exudates and rhizosphere microbiomes jointly determine temporal shifts in plant-soil feedbacks. Plant Cell Environ. 2023; doi: 10.1111/pce.14570.

54. Henderson D, Tello JS, Cayola L, Fuentes AF, Alvestegui B, Muchhala N, et al.. Testing the role of biotic interactions in shaping elevational diversity gradients: An ecological metabolomics approach. Ecology. 2025; doi: 10.1002/ecy.70069.

55. Bovay B, Descombes P, Chittaro Y, Glauser G, Nomoto H, Rasmann S. Adapting to change: Exploring the consequences of climate-induced host plant shifts in two specialist Lepidoptera species. Ecol Evol. 2024; doi: 10.1002/ece3.11596.

56. Sierra AM, Meléndez O, Bethancourt R, Bethancourt A, Rodríguez-Castro L, López CA, et al.. Leaf Endophytes Relationship with Host Metabolome Expression in Tropical Gymnosperms. J Chem Ecol. 2024; doi: 10.1007/s10886-024-01511-z.

57. Mleziva AD, Ngumbi EN. Comparative analysis of defensive secondary metabolites in wild teosinte and cultivated maize under flooding and herbivory stress. Physiol Plant. 2024; doi: 10.1111/ppl.14216.

58. Gallon ME, Muchoney ND, Smilanich AM. Viral Infection Induces Changes to the Metabolome, Immune Response and Development of a Generalist Insect Herbivore. J Chem Ecol. 2024; doi: 10.1007/s10886-024-01472-3.

59. Contreras-Cornejo HA, Schmoll M, Esquivel-Ayala BA, González-Esquivel CE, Rocha-Ramírez V, Larsen J. Mechanisms for plant growth promotion activated by Trichoderma in natural and managed terrestrial ecosystems. Microbiol Res. 2024; doi: 10.1016/j.micres.2024.127621.

60. Yasmin F, Cowie AE, Zerbe P. Understanding the chemical language mediating maize immunity and environmental adaptation. New Phytol. 2024; doi: 10.1111/nph.20000.

61. Ehlers BK, Berg MP, Staudt M, Holmstrup M, Glasius M, Ellers J, et al.. Plant Secondary Compounds in Soil and Their Role in Belowground Species Interactions. Trends Ecol Evol. Elsevier; 2020; doi: 10.1016/j.tree.2020.04.001.

62. Delory BM, Delaplace P, Fauconnier M-L, du Jardin P. Root-emitted volatile organic compounds: can they mediate belowground plant-plant interactions? Plant Soil. 2016; doi: 10.1007/s11104-016-2823-3.

63. Semchenko M, Barry KE, de Vries FT, Mommer L, Moora M, Maciá-Vicente JG. Deciphering the role of specialist and generalist plant–microbial interactions as drivers of plant–soil feedback. New Phytol. 2022; doi: 10.1111/nph.18118.

64. Ninkovic V, Markovic D, Rensing M. Plant volatiles as cues and signals in plant communication. Plant Cell Environ. 2021; doi: 10.1111/pce.13910.

65. Moore BD, Andrew RL, Külheim C, Foley WJ. Explaining intraspecific diversity in plant secondary metabolites in an ecological context. New Phytol. 2014; doi: 10.1111/nph.12526.

66. Bilas RD, Bretman A, Bennett T. Friends, neighbours and enemies: an overview of the communal and social biology of plants. Plant Cell Environ. 2021; doi: 10.1111/pce.13965.

67. Wang N-Q, Kong C-H, Wang P, Meiners SJ. Root exudate signals in plant–plant interactions. Plant Cell Environ. 2021; doi: 10.1111/pce.13892.

68. Rutz A, Sorokina M, Galgonek J, Mietchen D, Willighagen E, Gaudry A, et al.. The LOTUS initiative for open knowledge management in natural products research. Donoso DA, Akhmanova A, Tapley Hoyt C, editors. eLife. eLife Sciences Publications, Ltd; 2022; doi: 10.7554/eLife.70780.

69. Gaudry A, Pagni M, Mehl F, Moretti S, Quiros-Guerrero L-M, Cappelletti L, et al.. A Sample-Centric and Knowledge-Driven Computational Framework for Natural Products Drug Discovery. ACS Cent Sci. American Chemical Society; 2024; doi: 10.1021/acscentsci.3c00800.

70. Kim S, Chen J, Cheng T, Gindulyte A, He J, He S, et al.. PubChem 2025 update. Nucleic Acids Res. 2025; doi: 10.1093/nar/gkae1059.

71. Zdrazil B, Felix E, Hunter F, Manners EJ, Blackshaw J, Corbett S, et al.. The ChEMBL Database in 2023: a drug discovery platform spanning multiple bioactivity data types and time periods. Nucleic Acids Res. 2024; doi: 10.1093/nar/gkad1004.

72. Afendi FM, Okada T, Yamazaki M, Hirai-Morita A, Nakamura Y, Nakamura K, et al.. KNApSAcK Family Databases: Integrated Metabolite–Plant Species Databases for Multifaceted Plant Research. Plant Cell Physiol. 2012; doi: 10.1093/pcp/pcr165.

73. The Earth Metabolome Initiative. https://www.earthmetabolome.org/ (2023). Accessed 2026 Feb 14.

74. The Digital Botanical Gardens Initiative. https://digital-botanical-gardens-initiative.github.io/dbgi-green-paper/ (2022). Accessed 2025 June 11.

75. Wilkinson MD, Dumontier M, Aalbersberg IjJ, Appleton G, Axton M, Baak A, et al.. The FAIR Guiding Principles for scientific data management and stewardship. Sci Data. Nature Publishing Group; 2016; doi: 10.1038/sdata.2016.18.

76. Poelen JH, Simons JD, Mungall CJ. Global biotic interactions: An open infrastructure to share and analyze species-interaction datasets. Ecol Inform. 2014; doi: 10.1016/j.ecoinf.2014.08.005.

77. BiotXplorer. https://biotxplorer.sibils.org/. Accessed 2025 June 4.

78. Page R. Towards a biodiversity knowledge graph. Res Ideas Outcomes. Pensoft Publishers; 2016; doi: 10.3897/rio.2.e8767.

79. Kattge J, Díaz S, Lavorel S, Prentice IC, Leadley P, Bönisch G, et al.. TRY – a global database of plant traits. Glob Change Biol. 2011; doi: 10.1111/j.1365-2486.2011.02451.x.

80. Kattge J, Bönisch G, Díaz S, Lavorel S, Prentice IC, Leadley P, et al.. TRY plant trait database – enhanced coverage and open access. Glob Change Biol. 2020; doi: 10.1111/gcb.14904.

81. Wright IJ, Reich PB, Westoby M, Ackerly DD, Baruch Z, Bongers F, et al.. The worldwide leaf economics spectrum. Nature. Nature Publishing Group; 2004; doi: 10.1038/nature02403.

82. Nordt B, Hensen I, Bucher SF, Freiberg M, Primack RB, Stevens A-D, et al.. The PhenObs initiative: A standardised protocol for monitoring phenological responses to climate change using herbaceous plant species in botanical gardens. Funct Ecol. 2021; doi: 10.1111/1365-2435.13747.

83. Caspi R, Billington R, Keseler IM, Kothari A, Krummenacker M, Midford PE, et al.. The MetaCyc database of metabolic pathways and enzymes - a 2019 update. Nucleic Acids Res. 2020; doi: 10.1093/nar/gkz862.

84. Karp PD, Paley S, Caspi R, Kothari A, Krummenacker M, Midford PE, et al.. The EcoCyc Database (2023). EcoSal Plus. American Society for Microbiology; 2023; doi: 10.1128/ecosalplus.esp-0002-2023.

85. Karp PD, Billington R, Caspi R, Fulcher CA, Latendresse M, Kothari A, et al.. The BioCyc collection of microbial genomes and metabolic pathways. Brief Bioinform. 2019; doi: 10.1093/bib/bbx085.

86. Singh K, Maurya H, Singh P, Panda P, Behera AK, Jamal A, et al.. DISPEL: database for ascertaining the best medicinal plants to cure human diseases. Database. 2023; doi: 10.1093/database/baad073.

87. Youn J, Li F, Simmons G, Kim S, Tagkopoulos I. FoodAtlas: Automated Knowledge Extraction of Food and Chemicals from Literature. bioRxiv. 2024; doi: 10.1101/2024.05.16.594596.

88. The Earth Metabolome Initiative (EMI) ontology. https://w3id.org/emi. Accessed 2025 Aug 23.

89. The Earth Metabolome Initiative (EMI) ontology. https://www.earthmetabolome.org/earth_metabolome_ontology/. Accessed 2025 Aug 23.

90. Calvanese D, Lanti D, Mendes De Farias T, Mosca A, Xiao G. Accessing scientific data through knowledge graphs with Ontop. Patterns. 2021; doi: 10.1016/j.patter.2021.100346.

91. Calvanese D, Cogrel B, Komla-Ebri S, Kontchakov R, Lanti D, Rezk M, et al.. Ontop: Answering SPARQL queries over relational databases. Semantic Web. SAGE Publications; 2016; doi: 10.3233/SW-160217.

92. Xiao G, Lanti D, Kontchakov R, Komla-Ebri S, Güzel-Kalaycı E, Ding L, et al.. The Virtual Knowledge Graph System Ontop (Extended Abstract).

93. RDFlib. https://rdflib.dev/. Accessed 2025 June 6..

94. TRY Data Explorer. https://www.try-db.org/TryWeb/dp2.php. Accessed 2025 Aug 23.

95. Tandon D (2025, September 8). Plant traits data from TRY database (raw data for METRIN-KG) (Version v1). Zenodo. doi: 10.5281/zenodo.17079465.

96. Community GloBI (2025, January 13). Global Biotic Interactions: Interpreted Data Products (Version 0.8). Zenodo. doi: 10.5281/zenodo.14640564.

97. GloBI data. https://www.globalbioticinteractions.org/data. Accessed 2025 June 4.

98. GloBI GitHub repository. https://github.com/globalbioticinteractions. Accessed 2025 June 4.

99. GloBI datasets. https://www.globalbioticinteractions.org/datasets. Accessed 2025 June 4.

100. Bast H, Buchhold B. QLever: A Query Engine for Efficient SPARQL+Text Search. Proc 2017 ACM Conf Inf Knowl Manag. New York, NY, USA: Association for Computing Machinery; doi: 10.1145/3132847.3132921.

101. Qlever Wikidata SPARQL endpoint. https://qlever.dev/wikidata Accessed 2026 Feb 14.

102. Query to map wikidata identifiers to other taxonomies. https://qlever.dev/wikidata/S2gD0b Accessed 2026 Feb 14.

103. Query to retrieve lineage from Wikidata identifiers. https://qlever.dev/wikidata/66ksWA Accessed 2026 Feb 14.

104. Tandon D, Mendes de Farias T, Allard P-M, Defossez E. (2026, February 16). METRIN-KG Data (Version v7). Zenodo. doi: 10.5281/zenodo.19485732

105. Uber-Anatomy Ontology. https://purl.obolibrary.org/obo/uberon.owl. Accessed 2025 May 28.

106. Plant Ontology. https://purl.obolibrary.org/obo/po.owl. Accessed 2025 May 28.

107. Environment Ontology. https://purl.obolibrary.org/obo/envo.owl. Accessed 2025 May 28.

108. Gene Ontology. https://purl.obolibrary.org/obo/go.owl. Accessed 2025 May 28.

109. Phenotype and Trait Ontology. https://purl.obolibrary.org/obo/pato.owl. Accessed 2025 May 28.

110. METRIN-KG ontology matching at main · earth-metabolome-initiative/metrin-kg. GitHub. https://github.com/earth-metabolome-initiative/metrin-kg/tree/main/src/ontology_matching. Accessed 2026 Feb 14.

111. Lamy J-B. Owlready: Ontology-oriented programming in Python with automatic classification and high level constructs for biomedical ontologies. Artif Intell Med. 2017; doi: 10.1016/j.artmed.2017.07.002.

112. Reimers N, Gurevych I. Sentence-BERT: Sentence Embeddings using Siamese BERT-Networks. arXiv. 2019; doi: 10.48550/arXiv.1908.10084.

113. QUDT units vocabulary. https://qudt.org/doc/2025/01/DOC_VOCAB-UNITS-ALL.html. Accessed 2025 Aug 23.

114. FAIRsharing Team. FAIRsharing record for: Quantities, Units, Dimensions and Types. FAIRsharing (2015); doi: 10.25504/FAIRSHARING.D3PQW7.

115. W3C. https://www.w3.org/. Accessed 2025 Aug 23.

116. Janowicz K, Haller A, Cox SJD, Le Phuoc D, Lefrançois M. SOSA: A lightweight ontology for sensors, observations, samples, and actuators. J Web Semant. 2019; doi: 10.1016/j.websem.2018.06.003.

117. Advancing Geospatial Standards and Technology | OGC. Open Geospatial Consort. https://www.ogc.org/. Accessed 2025 June 4.

118. OBO Foundry. https://obofoundry.org/ontology/ro.html. Accessed 2025 Aug 23.

119. SKOS Simple Knowledge Organization System Namespace Document - HTML Variant, 18 August 2009 Recommendation Edition. https://www.w3.org/2009/08/skos-reference/skos.html. Accessed 2025 June 4.

120. W3C Semantic Web Interest Group: Basic Geo (WGS84 lat/long) Vocabulary. https://www.w3.org/2003/01/geo/. Accessed 2025 June 4.

121. The ENPKG RDF vocabulary. https://enpkg.commons-lab.org/doc/index.html. Accessed 2025 Aug 23.

122. Earth Metabolome Ontology GitHub repository. https://github.com/earth-metabolome-initiative/earth_metabolome_ontology. Accessed 2026 Feb 15.

123. Allard P-M, Gaudry A. (2024, March 17). Input and enriched files for the pf1600 dataset - ENPKG (Version 1.0). Zenodo. doi: 10.5281/zenodo.10827917.

124. Allard P-M, Gaudry A, Quirós-Guerrero L-M, Rutz A, Dounoue-Kubo M, Walker TWN, et al.. Open and reusable annotated mass spectrometry dataset of a chemodiverse collection of 1,600 plant extracts. GigaScience. 2023; doi: 10.1093/gigascience/giac124.

125. MySQL :: MySQL 8.2 Release Notes. https://dev.mysql.com/doc/relnotes/mysql/8.2/en/. Accessed 2025 June 6.

126. Earth Metabolome Initiative Ontology Ontop Mapping. The Earth Metabolome Initiative. https://github.com/earth-metabolome-initiative/earth_metabolome_ontology/blob/main/ontop_config/emi-v1/emi-v1.obda. Accessed 2025 Aug 23.

127. Krech D, Grimnes GAa, Higgins G, Hees J, Aucamp I, Lindström N, et al.. (2023, August 1). RDFLib (Version 7.0.0). Zenodo. doi: 10.5281/zenodo.8206632.

128. METRIN-KG GitHub repository case studies. The Earth Metabolome Initiative. https://github.com/earth-metabolome-initiative/metrin-kg. Accessed 2026 Feb 15.

129. Qlever-control GitHub repository. University of Freiburg: Algorithms and Data Structures Group. https://github.com/qlever-dev/qlever-control. Accessed 2026 Feb 15.

130. METRIN-KG SPARQL endpoint. https://kg.earthmetabolome.org/metrin/. Accessed 2026 Feb 15.

131. METRIN-KG Qlever SPARQL endpoint API. https://kg.earthmetabolome.org/metrin/api. Accessed 2026 Feb 15.

132. Programmatic access to METRIN-KG SPARQL endpoint. GitHub. https://github.com/earth-metabolome-initiative/metrin-kg/wiki/How-to-programatically-access-METRIN%E2%80%90KG%27s-SPARQL-endpoint%3F. Accessed 2026 Feb 15.

133. METRIN-KG SPARQL underlying endpoint. https://qlever.earthmetabolome.org/metrin-kg/. Accessed 2026 Feb 15.

134. Bast H, Kalmbach J, Klumpp T, Kramer F, Schnelle N. Efficient SPARQL Autocompletion via SPARQL. arXiv. 2021; doi: 10.48550/arXiv.2104.14595.

135. Qlever-ui GitHub repository. University of Freiburg: Algorithms and Data Structures Group. https://github.com/qlever-dev/qlever-ui. Accessed 2026 Feb 15.

136. Fork of Qlever-ui GitHub repository. The Earth Metabolome Initiative. https://github.com/earth-metabolome-initiative/qlever-ui. Accessed 2026 Feb 15.

137. METRIN-KG metrics-1 query. https://kg.earthmetabolome.org/metrin/metrics_1. Accessed 2026 Feb 15.

138. GloBI dataset template data format. https://github.com/globalbioticinteractions/template-dataset#data-format-and-dictionary. Accessed 2025 June 4.

139. METRIN-KG metrics-2 query. https://kg.earthmetabolome.org/metrin/metrics_2. Accessed 2026 Feb 15.

140. METRIN-KG metrics-3 query. https://kg.earthmetabolome.org/metrin/metrics_3. Accessed 2026 Feb 15.

141. METRIN-KG example query-11. https://kg.earthmetabolome.org/metrin/11. Accessed 2026 Feb 15.

142. METRIN-KG example query-11 versioned. GitHub. https://kg.earthmetabolome.org/metrin/11/v/53581b1 Accessed 2026 Feb 15.

143. METRIN-KG example query-12. https://kg.earthmetabolome.org/metrin/12. Accessed 2026 Feb 15.

144. METRIN-KG example query-12 versioned. GitHub. https://kg.earthmetabolome.org/metrin/12/v/53581b1. Accessed 2026 Feb 15.

145. METRIN-KG example query-13. https://kg.earthmetabolome.org/metrin/13. Accessed 2026 Feb 15.

146. METRIN-KG example query-13 versioned. GitHub. https://kg.earthmetabolome.org/metrin/13/v/53581b1. Accessed 2026 Feb 15.

147. METRIN-KG example query-14. https://kg.earthmetabolome.org/metrin/14. Accessed 2026 Feb 15.

148. METRIN-KG example query-14 versioned. GitHub. https://kg.earthmetabolome.org/metrin/14/v/53581b1. Accessed 2026 Feb 15.

149. METRIN-KG example query-20. https://kg.earthmetabolome.org/metrin/20. Accessed 2026 Feb 15.

150. METRIN-KG example query-20 versioned. GitHub. https://kg.earthmetabolome.org/metrin/20/v/53581b1. Accessed 2026 Feb 15.

151. METRIN-KG example query-18. https://kg.earthmetabolome.org/metrin/18. Accessed 2026 Feb 15.

152. METRIN-KG example query-18 versioned. GitHub. https://kg.earthmetabolome.org/metrin/18/v/53581b1. Accessed 2026 Feb 15.

153. METRIN-KG example query-22. https://kg.earthmetabolome.org/metrin/22. Accessed 2022 Feb 15.

154. METRIN-KG example query-22 versioned. GitHub. https://kg.earthmetabolome.org/metrin/22/v/53581b1. Accessed 2026 Feb 15.

155. METRIN-KG example query-23. https://kg.earthmetabolome.org/metrin/23. Accessed 2026 Feb 15.

156. METRIN-KG example query-23 versioned. https://kg.earthmetabolome.org/metrin/23/v/53581b1. Accessed 2026 Feb 15.

157. METRIN-KG example query-16. https://kg.earthmetabolome.org/metrin/16. Accessed 2026 Feb 15.

158. METRIN-KG example query-16 versioned. GitHub. https://kg.earthmetabolome.org/metrin/16/v/53581b1. Accessed 2026 Feb 15.

159. METRIN-KG example query-17. https://kg.earthmetabolome.org/metrin/17. Accessed 2026 Feb 15.

160. METRIN-KG example query-17 versioned. GitHub. https://kg.earthmetabolome.org/metrin/17/v/53581b1. Accessed 2026 Feb 15.

161. METRIN-KG example query-19. https://kg.earthmetabolome.org/metrin/19. Accessed 2026 Feb 15.

162. METRIN-KG example query-19 versioned. GitHub. https://kg.earthmetabolome.org/metrin/19/v/53581b1. Accessed 2026 Feb 15.

163. METRIN-KG example query-21. https://kg.earthmetabolome.org/metrin/21. Accessed 2026 Feb 15.

164. METRIN-KG example query-21 versioned. GitHub. https://kg.earthmetabolome.org/metrin/21/v/53581b1. Accessed 2025 Aug 23.

165. Bolleman J, Emonet V, Altenhoff A, Bairoch A, Blatter M-C, Bridge A, et al.. A large collection of bioinformatics question-query pairs over federated knowledge graphs: methodology and applications. arXiv; 2024; doi: 10.48550/arXiv.2410.06010

166. Emonet V, Sima A-C, Farias TM de. A user-friendly SPARQL query editor powered by lightweight metadata. arXiv; 2025; doi: 10.48550/arXiv.2503.02688

167. Contribute queries to METRIN-KG. https://github.com/earth-metabolome-initiative/metrin-kg/wiki/Contribute-queries-to-METRIN%E2%80%90KG. Accessed 2025 Aug 23.

168. Jupp S, Malone J, Bolleman J, Brandizi M, Davies M, Garcia L, et al.. The EBI RDF platform: linked open data for the life sciences. Bioinformatics. 2014; doi: 10.1093/bioinformatics/btt765.

169. Emonet V, Bolleman J, Duvaud S, Farias TM de, Sima AC. LLM-based SPARQL Query Generation from Natural Language over Federated Knowledge Graphs. arXiv. 2025; doi: 10.48550/arXiv.2410.06062.

170. ExpasyGPT for METRIN-KG. GitHub. https://github.com/earth-metabolome-initiative/metrin-kg/wiki/ExpasyGPT-for-METRIN%E2%80%90KG Accessed 2025 Aug 23.

171. Calleja JA, Domènech G, Sáez L, Lara F, Garilleti R, Albertos B. Extinction risk of threatened and non-threatened mosses: Reproductive and ecological patterns. Glob Ecol Conserv. 2022; doi: 10.1016/j.gecco.2022.e02254.

172. Gürlek S, Araújo AC, Brummitt N. Predicting the Threat Status of Mosses Using Functional Traits. Plants. Multidisciplinary Digital Publishing Institute; 2024; doi: 10.3390/plants13152019.

173. Junaedi DI, Nasution T, Putri DM, Iryadi R, Lestari R, Kurniawan V, et al.. Threatened exotic species of botanical gardens: Application of trait-based naturalized species risk scoring assessment. South Afr J Bot. 2023; doi: 10.1016/j.sajb.2022.11.046.

174. Álvarez-Yépiz JC, Búrquez A, Martínez-Yrízar A, Dovciak M. A trait-based approach to the conservation of threatened plant species. Oryx. 2019; doi: 10.1017/S003060531800087X.

175. Hill JL, Grisnik M, Hanscom RJ, Sukumaran J, Higham TE, Clark RW. The past, present, and future of predator–prey interactions in a warming world: Using species distribution modeling to forecast ectotherm–endotherm niche overlap. Ecol Evol. 2024; doi: 10.1002/ece3.11067.

176. Stelling-Wood TP, Poore AGB, Hughes AR, Everett JD, Gribben PE. Habitat traits and predation interact to drive abundance and body size patterns in associated fauna. Ecol Evol. 2023; doi: 10.1002/ece3.10771.

177. Ray K, Basak SK, Giri CK, Kotal HN, Mandal A, Chatterjee K, et al.. Ecological restoration at pilot-scale employing site-specific rationales for small-patch degraded mangroves in Indian Sundarbans. Sci Rep. Nature Publishing Group; 2024; doi: 10.1038/s41598-024-63281-8.

178. Mendes SB, Nogales M, Vargas P, Olesen JM, Marrero P, Romero J, et al.. Climb forest, climb: diverse disperser communities are key to assist plants tracking climate change on altitudinal gradients. New Phytol. 2025; doi: 10.1111/nph.20300.

179. Flickinger HD, Dukes JS. A Review of Theory: Comparing Invasion Ecology and Climate Change-Induced Range Shifting. Glob Change Biol. 2024; doi: 10.1111/gcb.17612.

180. Wang X, Cao Y, Jin Y, Sun L, Tang F, Dong L. Ecophysiological Trade-Off Strategies of Three Gramineous Crops in Response to Root Extracts of Phytolacca americana. Plants. Multidisciplinary Digital Publishing Institute; 2024; doi: 10.3390/plants13213026.

181. De La Peña R, Sattely ES. Rerouting plant terpene biosynthesis enables momilactone pathway elucidation. Nat Chem Biol. Nature Publishing Group; 2021; doi: 10.1038/s41589-020-00669-3.

182. Knoch E, Kovács J, Deiber S, Tomita K, Shanmuganathan R, Serra Serra N, et al.. Transcriptional response of a target plant to benzoxazinoid and diterpene allelochemicals highlights commonalities in detoxification. BMC Plant Biol. 2022; doi: 10.1186/s12870-022-03780-w.

183. Lu X, Zhang J, Brown B, Li R, Rodríguez-Romero J, Berasategui A, et al.. Inferring Roles in Defense from Metabolic Allocation of Rice Diterpenoids. Plant Cell. 2018; doi: 10.1105/tpc.18.00205.

184. Zhou S, Zhang R, Wang Q, Zhu J, Zhou J, Sun Y, et al.. OsbHLH5 Synergically Regulates Phenolamide and Diterpenoid Phytoalexins Involved in the Defense of Rice Against Pathogens. Int J Mol Sci. Multidisciplinary Digital Publishing Institute; 2024; doi: 10.3390/ijms252212152.

185. Vela F, Anese S, Varela RM, Torres A, Molinillo JMG, Macías FA. Bioactive Diterpenes from the Brazilian Native Plant (Moquiniastrum pulchrum) and Their Application in Weed Control. Molecules. Multidisciplinary Digital Publishing Institute; 2021; doi: 10.3390/molecules26154632.

186. Maraia H, Charles-Dominique T, Tomlinson KW, Staver AC, Jorge LR, Gélin U, et al.. Substantial Insect Herbivory in a South African Savanna-Forest Mosaic: A Neglected Topic. Ecol Evol. 2024; doi: 10.1002/ece3.70466.

187. Zhao Y, Hu J, Zhou Z, Li L, Zhang X, He Y, et al.. Biofortified Rice Provides Rich Sakuranetin in Endosperm. Rice. 2024; doi: 10.1186/s12284-024-00697-w.

188. Khatibi SMH, Dimaano NG, Veliz E, Sundaresan V, Ali J. Exploring and exploiting the rice phytobiome to tackle climate change challenges. Plant Commun. Elsevier; 2024; doi: 10.1016/j.xplc.2024.101078.

189. Bian S, Li Z, Song S, Zhang X, Shang J, Wang W, et al.. Enhancing Crop Resilience: Insights from Labdane-Related Diterpenoid Phytoalexin Research in Rice (Oryza sativa L.). Curr Issues Mol Biol. Multidisciplinary Digital Publishing Institute; 2024; doi: 10.3390/cimb46090634.

190. Wang X, Li X, Zhao W, Hou X, Dong S. Current views of drought research: experimental methods, adaptation mechanisms and regulatory strategies. Front Plant Sci. Frontiers; 2024; doi: 10.3389/fpls.2024.1371895.

191. Yang H, Ji S, Wu D, Zhu M, Lv G. Effects of Root–Root Interactions on the Physiological Characteristics of Haloxylon ammodendron Seedlings. Plants. Multidisciplinary Digital Publishing Institute; 2024; doi: 10.3390/plants13050683.

192. Shi Y, He Y, Zheng Y, Liu X, Wang S, Xiong T, et al.. Characteristics of the phyllosphere microbial community and its relationship with major aroma precursors during the tobacco maturation process. Front Plant Sci. Frontiers; 2024; doi: 10.3389/fpls.2024.1346154.

193. Chain FE, Romano E, Leyton P, Paipa C, Catalán CAN, Fortuna M, et al.. Vibrational and structural study of onopordopicrin based on the FTIR spectrum and DFT calculations. Spectrochim Acta A Mol Biomol Spectrosc. 2015; doi: 10.1016/j.saa.2015.05.072.

194. Suzuki M, Iwasaki A, Suenaga K, Kato-Noguchi H. Phytotoxic activity of crop residues from Burdock and an active substance. J Environ Sci Health Part B. Taylor & Francis; 2019; doi: 10.1080/03601234.2019.1636600.

195. El Khatib N, Morel S, Hugon G, Rapior S, Carnac G, Saint N. Identification of a Sesquiterpene Lactone from Arctium lappa Leaves with Antioxidant Activity in Primary Human Muscle Cells. Molecules. Multidisciplinary Digital Publishing Institute; 2021; doi: 10.3390/molecules26051328.

196. Zhang J, Zheng Z-Q, Xu Q, Li Y, Gao K, Fang J. Onopordopicrin from the new genus Shangwua as a novel thioredoxin reductase inhibitor to induce oxidative stress-mediated tumor cell apoptosis. J Enzyme Inhib Med Chem. Taylor & Francis; 2021; doi: 10.1080/14756366.2021.1899169.

197. Maeta A, Okamoto Y, Ishikawa H, Matsunaga T, Takahashi K. Japanese Leaf Burdock Extract Inhibits Adipocyte Differentiation in 3T3-L1 Cells. Plant Foods Hum Nutr. 2025; doi: 10.1007/s11130-024-01257-9.

198. Jalloh AA, Khamis FM, Yusuf AA, Subramanian S, Mutyambai DM. Long-term push–pull cropping system shifts soil and maize-root microbiome diversity paving way to resilient farming system. BMC Microbiol. 2024; doi: 10.1186/s12866-024-03238-z.

199. Czarnobai De Jorge B, Koßmann A, Hummel HE, Gross J. Evaluation of a push-and-pull strategy using volatiles of host and non-host plants for the management of pear psyllids in organic farming. Front Plant Sci. Frontiers; 2024; doi: 10.3389/fpls.2024.1375495.

200. Khan ZR, Chiliswa P, Ampong-Nyarko K, Smart LE, Polaszek A, Wandera J, et al.. Utilisation of Wild Gramineous Plants for Management of Cereal Stemborers in Africa. Int J Trop Insect Sci. 1997; doi: 10.1017/S1742758400022268.

201. Midega CAO, Wasonga CJ, Hooper AM, Pickett JA, Khan ZR. Drought-tolerant Desmodium species effectively suppress parasitic striga weed and improve cereal grain yields in western Kenya. Crop Prot. 2017; doi: 10.1016/j.cropro.2017.03.018.

202. Khan ZR, Midega CAO, Amudavi DM, Hassanali A, Pickett JA. On-farm evaluation of the ‘push–pull’ technology for the control of stemborers and striga weed on maize in western Kenya. Field Crops Res. 2008; doi: 10.1016/j.fcr.2007.12.002.

203. Hooper AM, Caulfield JC, Hao B, Pickett JA, Midega CAO, Khan ZR. Isolation and identification of Desmodium root exudates from drought tolerant species used as intercrops against Striga hermonthica. Phytochemistry. 2015; doi: 10.1016/j.phytochem.2015.06.026.

204. Reich PB, Walters MB, Ellsworth DS. From tropics to tundra: Global convergence in plant functioning. Proc Natl Acad Sci. Proceedings of the National Academy of Sciences; 1997; doi: 10.1073/pnas.94.25.13730.

205. Tandon D, Mendes De Farias T, Allard P-M, Defossez E. (2026, February 18). METRIN-KG: A knowledge graph integrating plant metabolites, traits and biotic interactions (Version v1.0.2). Zenodo. doi: 10.5281/zenodo.18684960

206. Agronomy Ontology. https://purl.obolibrary.org/obo/agro.owl. Accessed 2025 May 28.

207. Ascomycete Phenotype Ontology. https://purl.obolibrary.org/obo/apo.owl. Accessed 2025 May 28.

208. Biological Spatial Ontology. https://purl.obolibrary.org/obo/bspo.owl. Accessed 2025 May 28.

209. BRENDA Tissue Ontology. https://purl.obolibrary.org/obo/bto.owl. Accessed 2025 May 28.

210. Cephalopod Ontology. https://purl.obolibrary.org/obo/ceph.owl. Accessed 2025 May 28.

211. Chemical Entities of Biological Interest. https://purl.obolibrary.org/obo/chebi.owl. Accessed 2025 May 28.

212. Cell Ontology. https://purl.obolibrary.org/obo/cl.owl. Accessed 2025 May 28.

213. Collembola Anatomy Ontology. https://purl.obolibrary.org/obo/clao.owl. Accessed 2025 May 28.

214. Dictyostelium Discoideum Anatomy. https://purl.obolibrary.org/obo/ddanat.owl. Accessed 2025 May 28.

215. Ecology Core Ontology. https://purl.obolibrary.org/obo/ecocore.owl. Accessed 2025 May 28.

216. Experimental Factor Ontology. https://www.ebi.ac.uk/efo/efo.owl. Accessed 2025 May 28.

217. Fungal Anatomy Ontology. https://purl.obolibrary.org/obo/fao.owl. Accessed 2025 May 28.

218. Flora Phenotype Ontology. https://purl.obolibrary.org/obo/flopo.owl. Accessed 2025 May 28.

219. Foundational Model of Anatomy. https://purl.obolibrary.org/obo/fma.owl. Accessed 2025 May 28.

220. Food Ontology. https://purl.obolibrary.org/obo/foodon.owl. Accessed 2025 May 28.

221. Ontology for Plant Gall Phenotypes. https://purl.obolibrary.org/obo/gallont.owl. Accessed 2025 May 28.

222. Hymenoptera Anatomy Ontology. https://purl.obolibrary.org/obo/hao.owl. Accessed 2025 May 28.

223. Infectious Disease Ontology. https://purl.obolibrary.org/obo/ido.owl. Accessed 2025 May 28.

224. Infectious Disease Ontology for Malaria. https://purl.obolibrary.org/obo/idomal.owl. Accessed 2025 May 28.

225. National Cancer Institute Thesaurus. https://purl.obolibrary.org/obo/ncit.owl. Accessed 2025 May 28.

226. Ontology for Biomedical Investigations. https://purl.obolibrary.org/obo/obi.owl. Accessed 2025 May 28.

227. Ontology for MIcroRNA Target Prediction. https://purl.obolibrary.org/obo/omit.owl. Accessed 2025 May 28.

228. Pathogen-Host Interaction Phenotype Ontology. https://purl.obolibrary.org/obo/phipo.owl. Accessed 2025 May 28.

229. Porifera Ontology. https://purl.obolibrary.org/obo/poro.owl. Accessed 2025 May 28.

230. Mosquito gross anatomy ontology. https://purl.obolibrary.org/obo/tgma.owl. Accessed 2025 May 28.

