## Supplementary Data for "METRIN-KG: A knowledge graph integrating plant metabolites, traits, and biotic interactions"

SPARQL queries to Qlever’s Wikidata SPARQL endpoint

1. Query [101] for mapping Wikidata identifiers to ones from 15 taxonomy databases.

```
`PREFIX wdt: <http://www.wikidata.org/prop/direct/>

PREFIX wd: <http://www.wikidata.org/entity/>

SELECT ?WdID ?eol ?gbif ?ncbi ?ott ?itis ?irmng ?col ?nbn ?worms ?bold ?plazi ?apni ?msw3 ?iNat ?eppo ?WdName WHERE {

    ?WdID wdt:P31 wd:Q16521;

            wdt:P225 ?WdName .

    OPTIONAL { ?WdID wdt:P9157 ?ott . }

    OPTIONAL { ?WdID wdt:P685 ?ncbi . }

    OPTIONAL { ?WdID wdt:P846 ?gbif . }

    OPTIONAL { ?WdID wdt:P830 ?eol . }

    OPTIONAL { ?WdID wdt:P815 ?itis . }

    OPTIONAL { ?WdID wdt:P5055 ?irmng . }

    OPTIONAL { ?WdID wdt:P10585 ?col . }

    OPTIONAL { ?WdID wdt:P3240 ?nbn . }

    OPTIONAL { ?WdID wdt:P850 ?worms . }

    OPTIONAL { ?WdID wdt:P3606 ?bold . }

    OPTIONAL { ?WdID wdt:P1992 ?plazi . }
```

```
    OPTIONAL { ?WdID wdt:P5984 ?apni . }

    OPTIONAL { ?WdID wdt:P959 ?msw3 . }

    OPTIONAL { ?WdID wdt:P3151 ?iNat . }

    OPTIONAL { ?WdID wdt:P3031 ?eppo . }

  }`
```

2. Query [102] for retrieving Wikidata identifiers and their lineage

```
`PREFIX wdt: <http://www.wikidata.org/prop/direct/>

PREFIX wd: <http://www.wikidata.org/entity/>

SELECT ?WdID ?WdName ?hTax ?hTaxName ?hTaxRank WHERE {

    ?WdID wdt:P31 wd:Q16521;

        wdt:P225 ?WdName ;

        wdt:P171* ?hTax .

    ?hTax wdt:P225 ?hTaxName ;

        wdt:P105 ?hTaxRank .

}`
```
